# Dual functions of *labial* resolve the Hox logic of chelicerate head segments

**DOI:** 10.1101/2022.11.29.518396

**Authors:** Guilherme Gainett, Benjamin C. Klementz, Pola O. Blaszczyk, Heather Bruce, Nipam Patel, Prashant P. Sharma

## Abstract

Despite an abundance of gene expression surveys, comparatively little is known about Hox gene function in Chelicerata, with emphasis on the Hox logic of the anterior prosomal segments, which bear the mouthparts. Previous investigations of individual paralogs of *labial* (*lab*) and *Deformed* (*Dfd*) in the spider *Parasteatoda tepidariorum* have shown that these play a role in tissue maintenance of the pedipalpal segment (*labial-1*) and in patterning the first walking leg identity (*Deformed-1*), respectively. However, broader extrapolations of these data points across chelicerates are hindered by the existence of duplicated copies of Hox genes in arachnopulmonates (e.g., spiders and scorpions), which have resulted from an ancient whole genome duplication event. Here, we investigated the function of single-copy orthologs of *lab* in the harvestman *Phalangium opilio*, an exemplar of a lineage that was not subject of this whole genome duplication. Embryonic RNAi against *lab* resulted in homeotic transformations of pedipalps to chelicerae, as well as reduction and fusion of the pedipalpal segment with adjacent segments. To test for combinatorial function, we performed double knockdown of *lab* and *Dfd*, which results in homeotic transformation of both pedipalps and first walking legs into cheliceral identity, whereas the second walking leg is transformed into a pedipalpal identity. Taken together, these results elucidate a model for the Hox logic of head segments in Chelicerata. To substantiate the validity of this model, we additionally performed expression surveys for duplicated copies of *lab* and *Dfd* in scorpions and horseshoe crabs, toward understanding the genetic basis of a heteronomous prosoma. We show that repetition of morphologically similar appendages is correlated with uniform expression levels of the Hox genes *lab* and *Dfd*, irrespective of the number of gene copies.

## Introduction

The study of body plan evolution has a rich history, punctuated by advances in comparative embryology, genomics, and functional approaches. Arthropod body plan evolution has served as a particularly fertile proving ground for numerous concepts in evolutionary developmental biology, such as tests of serial homology, cooption of regulatory networks, evolution of cis-regulatory elements, and developmental systems drift (e.g (Gompel et al. 2005; Jager et al. 2006; Oliver et al. 2014; Auman and Chipman 2018; Setton and Sharma 2018)). Among the central contributions is the mechanistic understanding of arthropod Hox genes, a family of conserved transcription factors that play a role in regionalizing the antero-posterior axis across Bilateria. Numerous expression surveys have highlighted conserved and labile aspects of Hox expression domains across the arthropod tree of life, and advances in functional toolkits for emerging model species have pinpointed both canonical and non-canonical roles of these genes in body plan patterning (e.g (Abzhanov and Kaufman 1999; Hughes and Kaufman 2002a; Brena et al. 2006; Mahfooz et al. 2007; Medved et al. 2015; Martin et al. 2016)). However, the availability of functional datasets for Hox genes is highly asymmetrical, with much of what is known about anterior Hox genes stemming from pancrustacean (and particularly, insect) datasets. Whereas fundamentals of Hox gene dynamics are well understood in pancrustacean models like the fruit fly *Drosophila melanogaster* and the amphipod *Parhyale hawaiensis*, the functions of their homologs remain poorly understood in Chelicerata (e.g., sea spiders, horseshoe crabs, arachnids), the sister group to the remaining arthropods.

The bauplan of Chelicerata—and specifically, of arachnids—generally consists of two tagmata, the anterior prosoma (which bears the mouthparts and walking legs) and the posterior opisthosoma (which bears modified appendages or may lack appendages altogether in different groups) (Sharma, Schwager, et al. 2014). The mouthparts of chelicerates typically consist of a pair of chelicerae, innervated by the deutocerebrum (middle region of the tripartite brain; (Damen et al. 1998; Telford and Thomas 1998); and a pair of pedipalps, innervated by the tritocerebrum (posterior region of the tripartite brain). Posterior to these are segments that bear four pairs of legs, which are variably modified across chelicerate diversity. Modifications of this general architecture are found in Pycnogonida (sea spiders), which lack an opisthosoma and may bear additional leg pairs (e.g., the ovigers; ten- and twelve-legged species in some genera) (Ballesteros et al. 2021). Additionally, Xiphosura (the horseshoe crabs) exhibit anatomically identical pedipalps and walking legs with respect to the number of podomeres (leg segments); only the last pair of walking legs (the pusher leg) exhibits a distinct morphology, owing to its larger size, leaf-like terminal ornamentation, and the presence of the flabellum (an exite; see discussion in (Bruce 2021)). Whereas the tritocerebral appendage of Xiphosura is sexually dimorphic (comparable to the pedipalps of spiders or the third walking legs of Ricinulei), in females, this appendage is indistinguishable from the walking legs in the three immediately posterior segments, barring minor differences in size. As a result, Xiphosura is variably described in the literature as having pedipalps and four pairs of legs, or as having no pedipalps and five pairs of legs.

Functional datasets informing the patterning of chelicerate prosomal segments have long remained fragmentary. In the spider *Parasteatoda tepidariorum*, it was previously shown that one copy of the Hox gene *labial* (termed *labial-1*; abbr. *lab-1*) is necessary for the maintenance of the pedipalpal and L1 segment; maternal RNAi against this gene resulted in the reduction or complete deletion of the tritocerebral segment (and occasionally, also the L1 segment; (Pechmann et al. 2015). The implied function of segment maintenance is contrary to the broadly understood canonical role of Hox genes as drivers of homeosis. By contrast, RNAi against a copy of *Deformed* (termed *Deformed-1*; abbr. *Dfd-1*) in the same species resulted in homeotic transformation of the first walking leg into a pedipalpal identity, as inferred from both the morphology of the transformed appendage and the increased expression of *lab 1* in the ectopic pedipalp (Pechmann et al. 2015). No functional outcomes were reported from RNAi against the paralogs of each gene (i.e., spider *lab-2* and *Dfd-2*). The duplicates of these Hox genes are attributable to a shared genome duplication in the common ancestor of Arachnopulmonata, a group of six orders that includes spiders, scorpions, and pseudoscorpions (Sharma, Kaluziak, et al. 2014; Schwager et al. 2017; Ontano et al. 2021). Genomic datasets have shown that nearly all 20 Hox duplicates have been retained across all arachnopulmonate lineages, paralleling the evolutionary history of the vertebrates (Gainett and Sharma 2020; Harper et al. 2021).

However, as these duplicates are restricted to a subset of arachnid orders, it is not clear how well their dynamics reflect the ancestral condition of single-copy patterning genes, particularly given evidence for subfunctionalization or neofunctionalization of new gene copies in arachnopulmonate exemplars (Sharma, Schwager, et al. 2014; Turetzek et al. 2015; Benton et al. 2016; Turetzek et al. 2017). In addition, their incidence and retention in models like *P. tepidariorum* raises the specter of functional redundancy and compensatory effects between paralogs in RNAi experiments. For this reason, functional datasets from chelicerate taxa that did not undergo genome duplication events are especially valuable for comparison. Accordingly, recently comparative studies have focused on the unprepossessing harvestman species *Phalangium opilio*, which exhibits an unduplicated genome and is amenable to gene silencing techniques(Sharma et al. 2012a; Sharma et al. 2012b; Sharma et al. 2013; Gainett et al. 2021). Functional datasets in the harvestman *Phalangium opilio* have shown that the single-copy *Dfd* ortholog is required for patterning the identity of the first two pairs of walking legs; knockdown of *Dfd* resulted in the homeotic transformation of both L1 and L2 into pedipalps. Whereas RNAi against *Sex combs reduced* (*Scr*) had no effect by itself, double knockdown of *Dfd* and *Scr* resulted in homeotic transformations of L1-L3 toward pedipalpal identity (Gainett et al. 2021). To date, no other functional data are available for the Hox genes of this species.

Here, we focused on discovering the functions of single-copy orthologs of anterior Hox genes in *P. opilio*, with the aim of understanding the Hox logic of the arachnid head segments. We show that knockdown of *lab* results in homeotic transformation of the pedipalp into cheliceral identity, whereas double knockdown of both *lab* and *Dfd* transforms both pedipalps and first walking legs into chelicerae. Having established a ground plan for the patterning of segmental identities in the harvestman, we explored the expression dynamics of *lab* and *Dfd* paralogs in a scorpion and a horseshoe crab, as points of comparison to spider and harvestman Hox genes. This survey substantiates a correlation between heterogeneity of Hox expression levels and the morphological disparity of prosomal appendages.

## Results

### *Expression of harvestman* labial

The single copy *Po-labial* (*Po-lab*) is expressed as a ring in the equator of the embryo prior to the formation of the germ band, and in dispersed cells towards the center of the encircled labial domain (Fig. 1a). In the stage where the antero-posterior axis forms and the anterior tagma (prosoma) becomes segmented, strong expression localizes to the entire tritocerebral and L1 segment, as land-marked by the segment polarity gene *Po-en* (Fig. 1b). *Po-lab* is also expressed in the posterior part of the L2 and L3 segments (Fig. 1b). During formation of limb buds, *Po-lab* is strongly expressed in the ectoderm of the pedipalp and L1 limb buds, and in the mesoderm of L2 and L3 nascent limb buds (Fig. 1c–d). With the sequential addition of posterior body segments, additional paired dots of expression appear in sequence adjacent to the ventral midline (Fig. 1e–g, arrows). A sharp anterior boundary of expression in the tritocerebral segment, strong expression in the tritocerebral and L1 appendages, and along the ventral ectoderm are maintained throughout the stages investigated. (Fig. 1c–g).

**Fig. 1.**
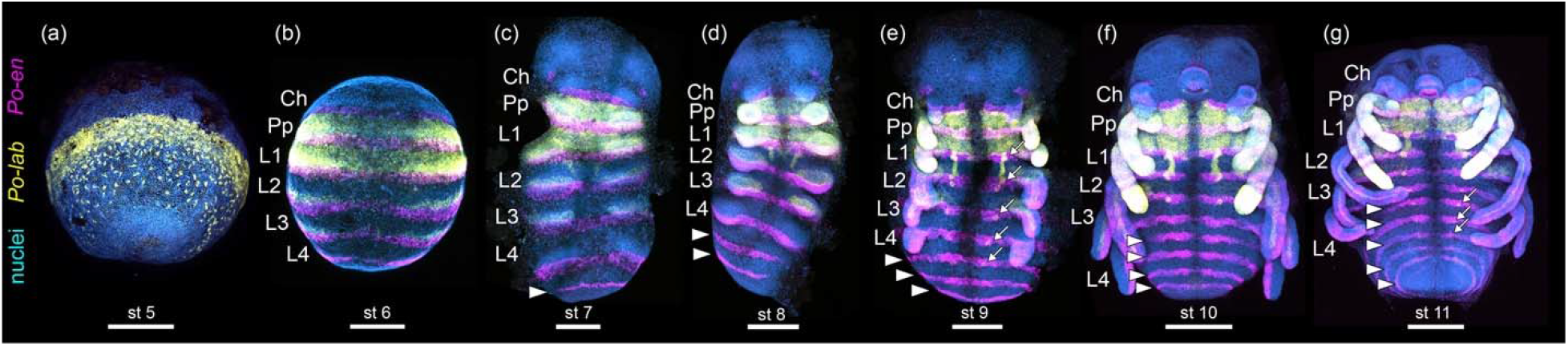
Wild type expression of *Popi-lab* and *Popi-en* across embryonic stages. Anterior at top. (a) Wholemount, lateral view of the germ disk. (b) Wholemount, ventral view. (c–g): Flatmount, ventral view. Nuclei in blue (Hoechst). *Popi-en*: magenta. *Popi-lab*: yellow. Arrowheads: *Popi-en* stripes marking the posterior of each segment. Arrows: dots of *Popi-lab* expression adjacent to the ventral midline. Ch: Chelicera; Pp: Pedipalp; L1–L4: L1–L4 legs. Scale bars: 200 µm.

### *Knockdown of* lab *results in homeotic pedipalp-to-chelicera transformations and defects in the tritocerebral segment*

To investigate the function of *Po-lab*, we conducted RNA interference (RNAi) via embryonic injections of double-stranded RNA (dsRNA). *Po-lab* RNAi hatchlings presented a spectrum of defects affecting the tritocerebral segment. The chelicera and leg identities were unaffected in hatchlings, whereas the pedipalps were transformed into cheliceral identity (n=35/74) (Fig. 2a-j; Supplementary File, Fig. S1, S2), as evidenced by a proximal podomere and a distal podomere with two claws (chela) (Fig 2b, g, l). Mild transformation consisted of partial truncation of the pedipalpal podomeres and unaffected claw (Fig. 2h, insets), whereas strong homeosis consisted of a tritocerebral appendage with proximal podomere and an ectopic claw in the distal podomere (Fig. 2h, insets). Homeosis in the embryonic appendages was evidenced by the branching and reduced length of the tritocerebral limb (Supplementary File, Fig. S1 a–b).

**Fig. 2.**
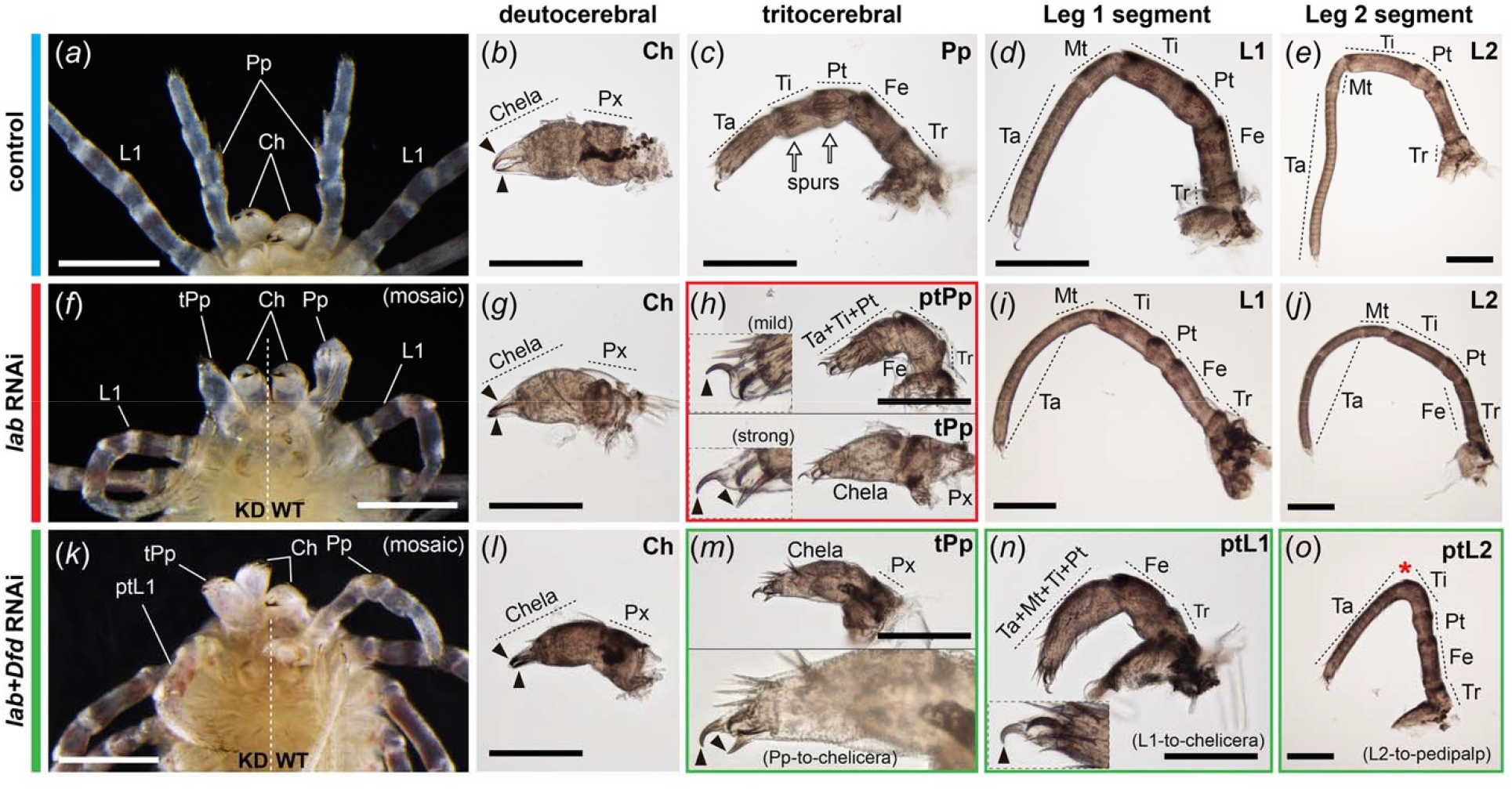
*Popi-lab* knockdown results in homeotic transformations. Brightfield images of hatchlings (postembryos) of Control (dH_2_O-injected) (a–e), Popi-lab RNAi (f–j), and *Popi-lab* + *Popi-Dfd* RNAi (k–o) experiments. Panels (a), (f) and (k) are ventral views of the anterior prosoma. Other panels are dissected appendages in lateral view with proximal at right. (b, g and l): Deutocerebral appendage (chelicera). (c, h, and m): Tritocerebral appendage. (d, i, and n): Appendage of the L1 segment. (e, j, and o): Appendage of the L2 segment. Note that, because the 1^st^ instar’s cuticle is already secreted and visible inside the postembryo cuticle, two sets of claws are discernible in some panels. Black arrowhead: claw; Red asterisk: missing metatarsus (Mt); White arrows: pedipalpal spurs; Ch: chelicera; Pp: pedipalp; tPp: transformed pedipalp; ptPp: partially transformed pedipalp; L1–L2: L1–L2 appendages; ptL1–ptL2: partially transformed L1–L2 appendages; Ta: tarsus; Mt: metatarsus; Ti: tibia; Pt: patella; Fe: femur; Tr: trochanter; Px: proximal cheliceral segment. Scale bars: 200 µm.

In addition to homeosis, a subset of *Po-lab* RNAi hatchlings (11%; n=4/35) exhibited defects in the segment boundaries of the tritocerebral segment, as evidenced by the fusion of the tritocerebral appendage to either the chelicera or to L1 (Fig. 3a–f; Supplementary File, Fig. S3). *Po-lab* RNAi embryos assayed for the segmentation marker *Po-engrailed* (*Po-en*) revealed a reduction in the tritocerebral and L1 segment relative to other prosomal segments, and juxtaposition of consecutive *Po-en* expression stripes (Supplementary File, Fig. S3). Taken together, these results suggest that in *P. opilio labial* is necessary for both maintenance of tritocerebral tissue and for the identity of the pedipalp.

**Fig. 3.**
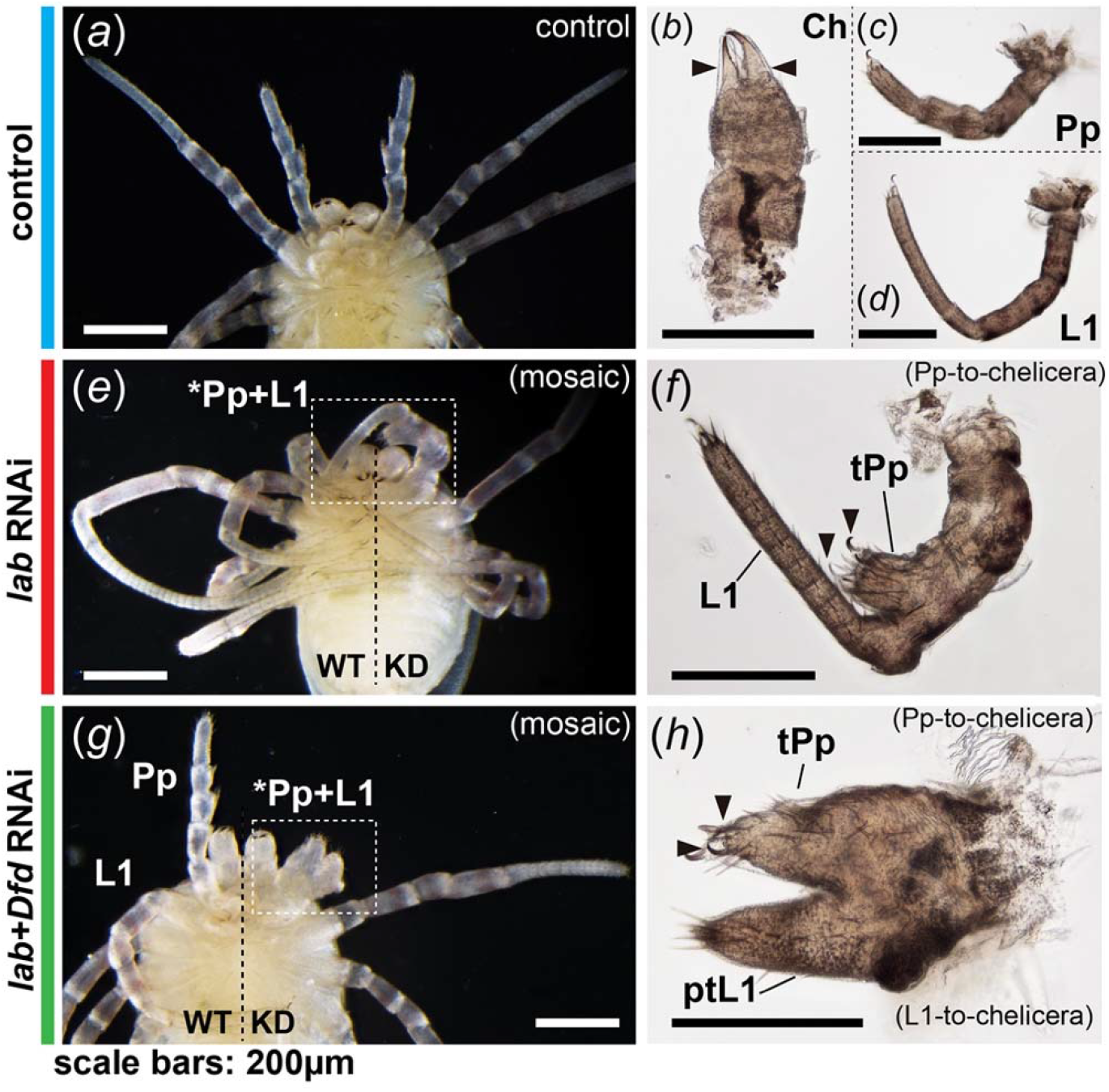
*Popi-lab* knockdown results in fusion of the pedipalp and L1 leg. Brightfield images of hatchlings (postembryos) of Control (dH_2_O-injected) (a–d), *Popi-lab* RNAi (e–f), and *Popi-lab* + *Popi-Dfd* RNAi (g–h) experiments. Phenotypes in g–h present both homeosis and fusion. Panels (a, e, and g) are ventral views of the anterior prosoma. Other panels are dissected appendages in lateral view. Mosaic individuals in (e) and (g) are affected on the right side of the picture. Black arrowhead: claw; Ch: chelicera; Pp: pedipalp; tPp: transformed pedipalp; L1: L1 appendage. Scale bars: 200 µm.

### *Double knockdown of* lab + Dfd *results in transformations of pedipalps and L1 legs into chelicerae*

While RNAi against *lab* results in transformation to cheliceral identity in the pedipalpal segment, we observed that the identity of L1 was not comparably affected, despite the presence and maintenance of *lab* transcript abundance in L1 territory. We inferred that that absence of function may be attributable to functional redundancy with another anteriorly expressed Hox gene such as *Dfd* and *proboscipedia* (*pb*), though the latter gene is only expressed in subdomains of the prosomal appendages (Sharma et al. 2012a). To test this possibility, we conducted RNAi against *proboscipedia* (*pb*) and double RNAi against *Po-lab* and *Po-Dfd*. Preliminary efforts with RNAi against *pb* resulted in no observable phenotype (data not shown); this outcome, together with the restriction of *pb* expression to only some of the tissue from the pedipalpal through L4 segment (Sharma et al. 2012a), suggested that *pb* may not act as a canonical Hox gene in *P. opilio*.

Double RNAi against *Po-lab* + *Po-Dfd* resulted in homeotic transformations (n=43/76) affecting tritocerebral, L1 and L2 segments (Fig. 2k–o). Fusions of the tritocerebral appendage with chelicera or L1 leg (n=9/76) were also observed, accompanied by homeosis in the pedipalp, L1 leg, or both appendages (Fig. 3f–g). A subset of the hatchlings (n=22/43) exhibit an additive phenotype, i.e., the combination of both pedipalp-to-chelicera (as in single *Po-lab* RNAi) and leg-to-pedipalp (as in single *Po-Dfd* RNAi (Gainett et al. 2021)), as evidenced by the truncation of the tritocerebral appendage, and loss of metatarsus in L1 leg. Notably, the effect in the legs in these cases is similar to the weak phenotypes of single *Po-Dfd* RNAi (Gainett et al. 2021), with a partial transformation of L1 and L2 legs into pedipalps. A smaller subset of hatchlings exhibited either single *Po-lab* RNAi phenotype (1/43) or single *Po-Dfd* RNAi (3/43) phenotype (Supplementary Fig. S2).

A new synergistic phenotype was also observed (17/43), in which both pedipalp-to chelicera and the L1 leg-to-chelicera homeosis occurred (Fig. 2k-o; Supplementary Fig. S2). Notably, we did not observe L2 leg-to-chelicera transformations, only partial L2 leg-to-pedipalp transformations, as evidenced by the lack of metatarsus (Fig. 2o), consistent with the absence of *lab* expression from the L2 segment.

These results are consistent with a canonical, combinatorial Hox model, wherein *lab* is required for patterning pedipalpal identity; *Dfd* for patterning the L1 and L2 legs (Gainett et al. 2021); and *Dfd* and *Scr* for patterning the L3 leg (Gainett et al. 2021).

### *Expression of* lab *and* Dfd *duplicates in a scorpion and a horseshoe crab*

A notable complexity in chelicerate Hox cluster evolution is the occurrence of paralogs incurrent from whole genome duplications, which parallels the history of the vertebrates (Wagner et al. 2003; Dehal and Boore 2005). Furthermore, the whole genome duplications in Arachnopulmonata (spiders, scorpions, and four other arachnid orders) and in Xiphosura (horseshoe crabs) are inferred to have occurred independently (Schwager et al. 2017; Shingate, Ravi, Prasad, Tay, Garg, et al. 2020; Ontano et al. 2021). Hox gene copies are of interest in comparative studies because they may undergo subfunctionalization or neofunctionalization, which in turn may correlate with diversification of body plans (Wagner et al. 2003; Sharma, Schwager, et al. 2014; Schwager et al. 2017). To test the validity of the Hox logic established herein, as well as obtain further insights into the fate of *lab* paralogs in Chelicerata, we investigated the expression patterns of *lab* paralogs in the scorpion *Centruroides sculpturatus* and in the horseshoe crab *Limulus polyphemus*.

The two *labial* paralogs of *C. sculpturatus* (*Cscu-lab 1, Cscu-lab 2*) have a clear anterior boundary of expression in the tritocerebral segment, and strong expression in the tritocerebral appendage (Fig. 4a–a’’). The two paralogs have complex patterns of expression on the ventral midline and developing nervous system, with several overlapping domains (Fig. 4a–a’’). The two *Dfd* paralogs (*Cscu-Dfd 1, Cscu-Dfd 2* have an anterior boundary of expression in the fourth head segment (L1) (Fig. 5 a–a’’). *Cscu-Dfd 1* is uniformly expressed in the L1–L4 legs and the ventral neuromeres (Fig. 5 a–a’’). *Cscu-Dfd 2* has largely overlapping expression with respect to its paralog; it differs from *Cscu-Dfd 1* in that it is more strongly expressed in the tips of the legs, it is slightly more anteriorly expressed in the L1 neuromere, and its expression in the opisthosoma is ubiquitous (Fig. 5 a–a’’). Therefore, the unique identity of the scorpion pedipalp with respect to the legs correlates with the strong expression of both *labial* paralogs on this appendage, whereas the identity of the four pairs of legs correlates with uniform expression of both *Deformed* paralogs in all legs.

**Fig. 4.**
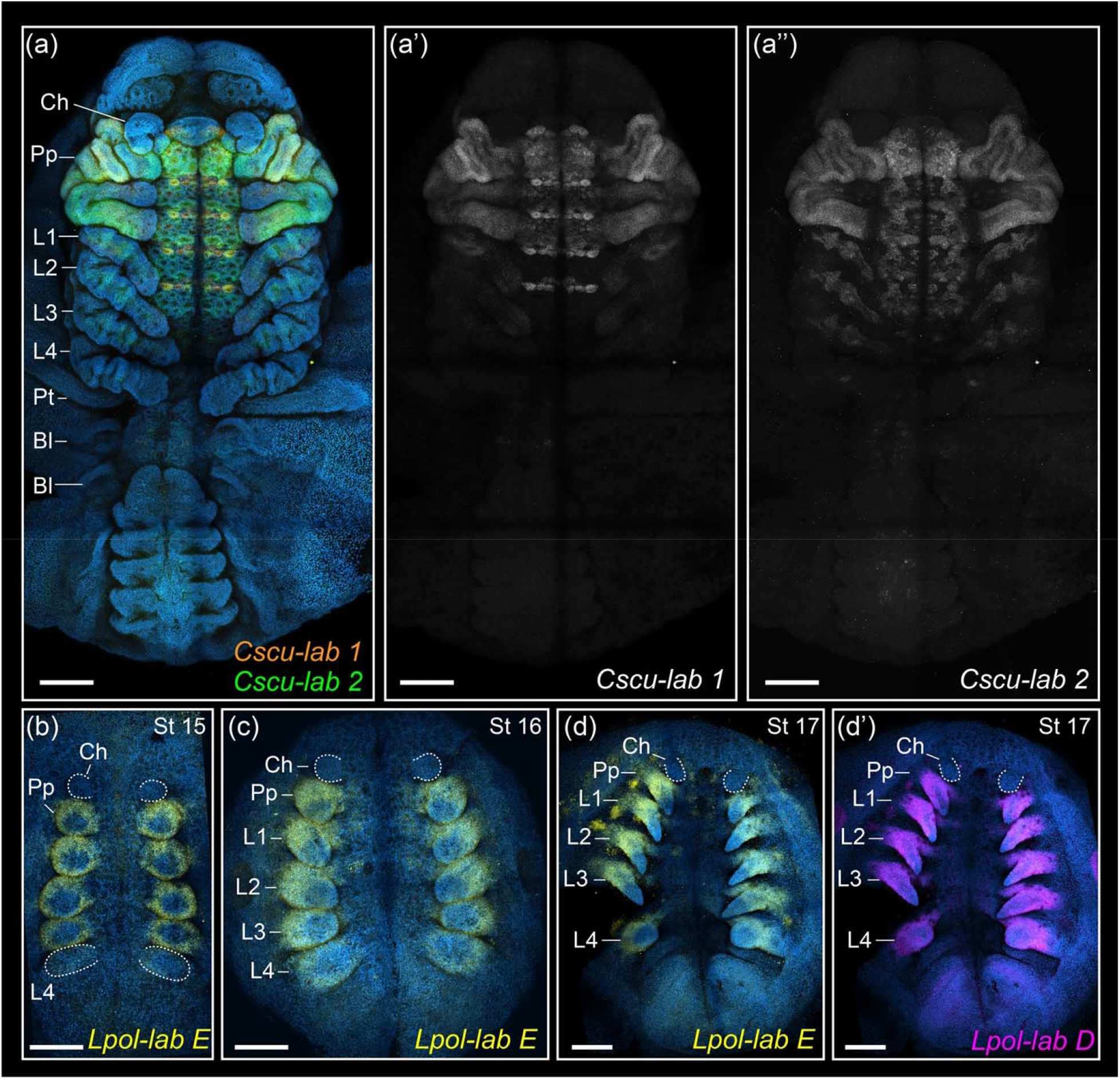
HCR in situ hybridization of *labial* paralogs in the scorpion *Centruroides sculpturatus* (a– a’’) and the horseshoe crab *Limulus polyphemus* (b–d’). Maximum intensity projections of flat mounted embryos in ventral view. Hoechst counter staining in blue. Same letters indicate same embryo (multiplexed). (a) Merged projections of *Cscu-lab1* (orange), *Cscu-lab 2* (green). Overlap of *lab* paralogs appears in yellow. (a’) Single channel projection of *Cscu-lab 1*. (a’’) Single channel projection of *Cscu-lab 2*. (b–d) *Lpol-lab E* expression (yellow). (d’’) *Lpol-lab D* expression (magenta). Ch: chelicera; Pp: pedipalp; L1–L4: L1–L4 legs. Scale bars: 200 µm.

**Fig. 5.**
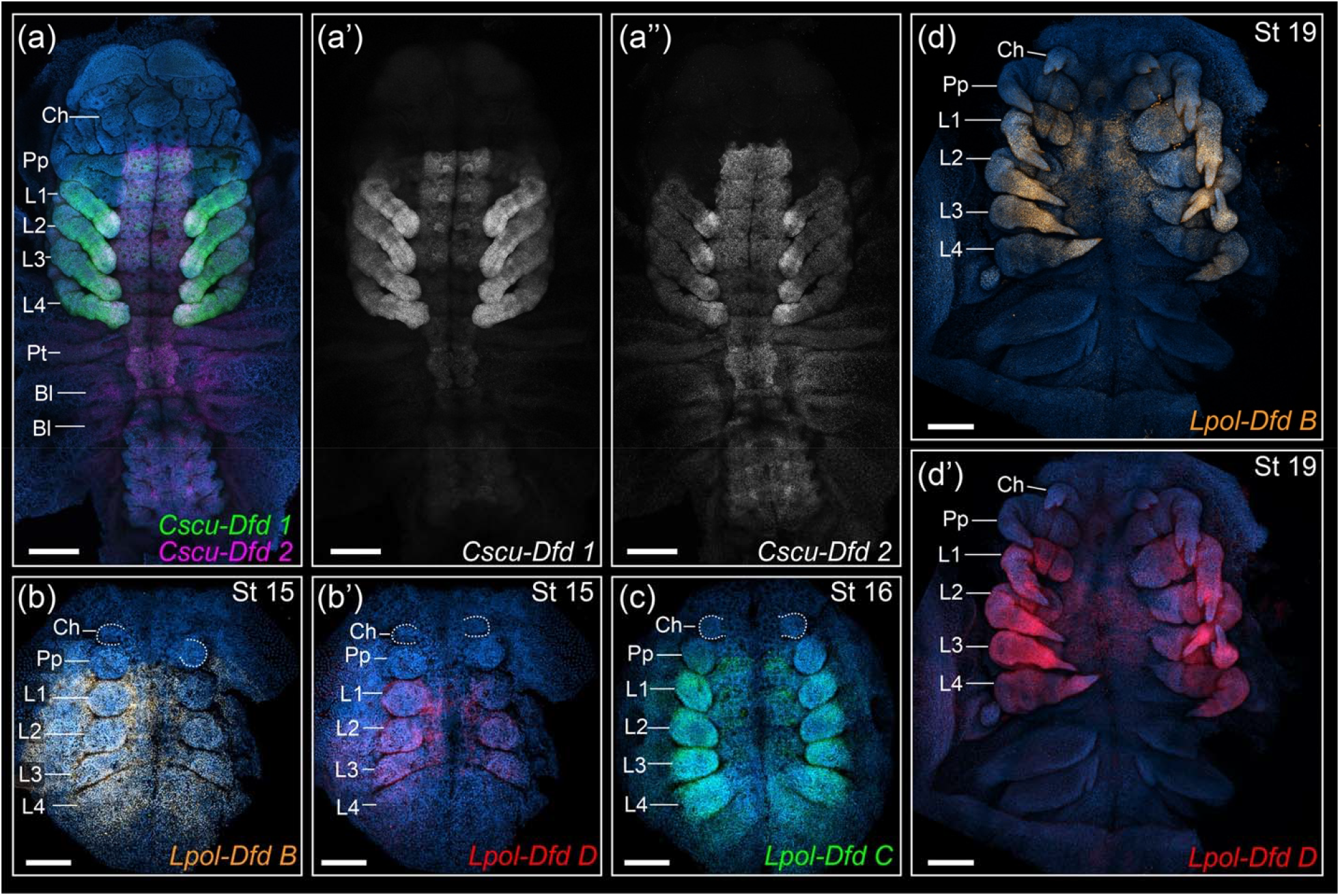
HCR in situ hybridization of *Deformed* paralogs in the scorpion *Centruroides sculpturatus* (a–a’’) and the horseshoe crab *Limulus polyphemus* (b–d’). Maximum intensity projections of flat mounted embryos in ventral view. Hoechst counter staining in blue. Same letters indicate same embryo (multiplexed). (a) Merged projections of *Cscu-Dfd 1* (green), *Cscu-Dfd 2* (magenta). (b) *Lpol-Dfd B* expression (orange). (b’) *Lpol-Dfd D* expression (red). (c) *Lpol-Dfd* C expression (green). (d) *Lpol-Dfd B* expression (orange). (d’) *Lpol-Dfd D* expression (red). Ch: chelicera; Pp: pedipalp; L1–L4: L1–L4 legs. Scale bars: 200 µm.

Four *labial* paralogs (*Lpol-lab A, Lpol-lab B, Lpol-lab D, Lpol-labE*) are present in the genome of *L. polyphemus* (five paralogs occur in *Tachypleus gigas* and *Carcinoscorpius rotundicauda*) (Kenny et al. 2015; Shingate, Ravi, Prasad, Tay, Garg, et al. 2020; Shingate, Ravi, Prasad, Tay, and Venkatesh 2020). *Lpol-lab E* has an anterior boundary of expression in the tritocerebral segment, being expressed in the tritocerebral appendage and posterior prosomal appendages throughout embryonic stages analysed (Fig. 4b–d). *Lpol-lab D* has a largely overlapping expression domain with respect to *Lpol-lab E*, with a sharp anterior boundary in the tritocerebral segment (Fig. 4 d’), but slightly more distally expressed in the appendages than *Lpol-lab E*. The short length of *Lpol-lab A* CDS (469bp) forestalled reliable detection of transcripts via HCR in situ hybridization. We did not analyze the putative homolog *Lpol-lab B*, given its unsual sequence and ambiguous annotation (Electronic Supplementary Methods).

Five *Dfd* paralogs occur in *L. polyphemus* (*Lpol-Dfd A–E*). *Lpol-Dfd B, Lpol-Dfd C*, and *Lpol-Dfd D* have a clear anterior boundary of expression in the L1 segment (Fig. 5 b–d’) and are expressed in the L1–L4 legs. The three paralogs have largely overlapping expression domains, with the exception that *Lpol-Dfd B* expression extends to the lateral margin of the germ band and more posteriorly into the opisthosoma at the germband stage (Fig. 5 b). We were not able to detect expression of *Lpol-Dfd A* and *Lpol-Dfd E*. Therefore, the shared morphology of the tritocerebral appendage and the subsequent prosomal appendage pairs (legs) correlates with a uniform expression of *labial* paralogs across all post-cheliceral prosomal appendages, and largely overlapping *lab* and *Dfd* paralog expression across the L1–L4 legs.

## Discussion

### labial *function in Arthropoda*

In the fruit fly *Drosophila melanogaster, lab* is expressed in the tritocerebral segment (the intercalary segment in insects, which is reduced and lacks appendages) during embryogenesis and is necessary for normal head involution (Merrill et al. 1989). However, there is no reported homeotic function during embryogenesis, which is also the case for a beetle and a milkweed bug (Angelini et al. 2005; Posnien and Bucher 2010; Schaeper et al. 2010). Homeosis in *lab* mutant flies is only detected in adults, as a shift from head-to-thorax identity of the bristles. This homeosis contrasts with the typical transformation towards anterior identity upon disruption of Hox genes.

By contrast, the tritocerebral segment of non-insect (and non-myriapod) arthropods (i.e., Chelicerata and the crustaceans) bears a pair of appendages that express *lab* (Telford and Thomas 1998; Abzhanov and Kaufman 1999; Jager et al. 2006; Sharma et al. 2012a; Serano et al. 2016). The only functional data point available for these groups, in the spider *Parasteatoda tepidariorum*, demonstrated a role in tritocerebral (and in some cases, L1) tissue maintenance, similarly to insects, but no homeotic function in tritocerebral appendage identity upon *lab-1* knockdown (Pechmann et al. 2015). In that work, knockdown of *Dfd-1* was also shown to result in homeotic transformation of L1 to pedipalps, with the ectopic pedipalps strongly expressing *lab-1* (Pechmann et al. 2015). This result is suggestive of the possibility that *lab* may play a role as a pedipalpal determinant, but the evidence is indirect, as homeosis induces transformation of the entire pedipalpal program and its associated gene expression patterns.

On the other hand, the spectrum of embryonic defects in *P. opilio* upon *lab* RNAi suggests a dual role for this gene, both in tritocerebral and L1 segment maintenance (segmental fusions in the phenotypic spectrum) and in canonical patterning of segmental identity (homeotic transformations of pedipalp to chelicera identity). These data reconcile the gap between the unexpected loss of the tritocerebral segment in spiders upon RNAi against *lab-1* and the previously unknown identity of the Hox gene that functions as the pedipalpal selector. It is possible that the *lab-1* segmentation phenotype in the spider reflects the severe end of the phenotypic spectrum, as a consequence of the mode and timing of delivery (maternal RNAi). This potential explanation could be tested via embryonic RNAi against *lab-1* in *P. tepidariorum*, with the prediction that lower concentrations of dsRNA injected at a later point in embryogenesis may elicit the homeotic phenotype.

More broadly, the significance of the *P. opilio lab* RNAi phenotype is that it represents the only known case of a homeotic function for *lab* aside from *D. melanogaster*. Given the phylogenetic relationship of these two species (a chelicerate and a hexapod), our results suggest that *lab* may play a role as a conserved tritocerebral selector across Arthropoda.

### The relationship between diminution of Hox expression and segmental deletions

The observation that severe loss-of-function phenotypes of *lab* homologs exhibit segment loss and/or fusion in two chelicerates (the spider *P. tepidariorum* (Pechmann et al. 2015) and the harvestman *P. opilio*) substantiates the intriguing possibility that loss or reduction of a Hox gene may be a potential mechanism for some segmental deletions across Panarthropoda. This inference is contrary to the archetypal role of Hox genes as selector genes that confer segmental identity. Deletion of Hox genes from the Hox cluster has been linked to segmental reduction in multiple panarthropod taxa, such as acariform mites (loss of segmentation posterior to the second opisthosomal segment, in tandem with loss of *abdominal-A* (Grbić et al. 2011; Barnett and Thomas 2013)) and tardigrades (loss of a large intermediate region of the antero-posterior axis, in tandem with multiple Hox genes (Smith et al. 2016)). Similarly, in sea spiders, the absence of a segmented opisthosoma has been tentatively linked to atypical patterns of sequence evolution in *abdominal-A* (Manuel et al. 2006), but expression patterns of posterior Hox genes are entirely missing in this group (Jager et al. 2006). The mechanism of Hox gene degradation and loss was thought to be released selection in the wake of segmental deletions, resulting in accumulation of mutations and eventual loss of nonfunctional genes associated with the deleted regions—Hox gene loss was understood to reflect the consequence of segmental loss, and not the cause (Barnett and Thomas 2013; Smith et al. 2016).

Nevertheless, it has been recently reported that diminished *Ubx* and *abd-A* expression in an ant results in posterior body truncations, which has been linked to a re-wiring of these genes to a lineage-specific function in early germ cell patterning (Rafiqi et al. 2020). In addition, several members of the homeobox gene family are classically known to be key to segmentation across Arthropoda, such as *even-skipped* and *caudal* (Hughes and Kaufman 2002b; Mito et al. 2007; Schönauer et al. 2016). Among these are the *Hox3* paralog *bicoid* in cyclorrhaphan flies and *fushi tarazu* (*ftz*) in a subset of insects. Indeed, in Arthropoda, evidence for a canonical Hox function is entirely absent for *Hox3* and greatly limited for *ftz*. Our results support the possibility that diminution or loss of *lab* expression may underlie the independent origins of the appendage-free and highly reduced intercalary segment of hexapods and myriapods, as previously articulated by Pechmann et al. (Pechmann et al. 2015). Given the absence of advanced functional toolkits for either the spider or the harvestman, tests of this hypothesis could capitalize upon the availability of genome editing tools in the amphipod crustacean *Parhyale hawaiensis* (Martin et al. 2016; Serano et al. 2016), with the goal of understanding how fine-tuning of *lab* expression regulates the transition between a homeotic transformation and a segmental deletion. More broadly, gene silencing tools developed in the tardigrade model *Hypsibius exemplaris* could be leveraged to test whether a non-canonical role of *lab* as a segmentation gene is an ancestral feature of Panarthropoda (Tenlen et al. 2013; Smith et al. 2016).

### Hox logic and evolution of the chelicerate prosoma

Leg identity in arachnids has been previously shown to require the expression of *Dfd*, as knockdown of *Dfd* results in leg-to-pedipalp homeotic transformation in the daddy-longlegs and in a spider (Pechmann et al. 2015; Gainett et al. 2021), with a redundant role of *Scr* in the case of L3 (Gainett et al. 2021). Similarly, it was previously shown that *homothorax* is required in the absence of Hox input for cheliceral identity (Sharma et al. 2015). The RNAi experiments with *lab* (and both *lab* and *Dfd*) presented herein thus enable the completion of a model for anterior prosomal fate specification in the harvestman (Fig. 6), wherein cheliceral identity is specified by *homothorax* in the absence of Hox input; pedipalpal identity requires high levels of *lab* and low levels of *Dfd*; and leg identity requires *Dfd* expression, with a redundant role of *Scr* at least in the case of the L3 appendage. The genetic basis for L4 identity remains unknown, but is likely attributable to *fushi tarazu*, which is strongly expressed in this territory in the manner of a canonical Hox gene (Sharma et al. 2012a).

**Fig. 6.**
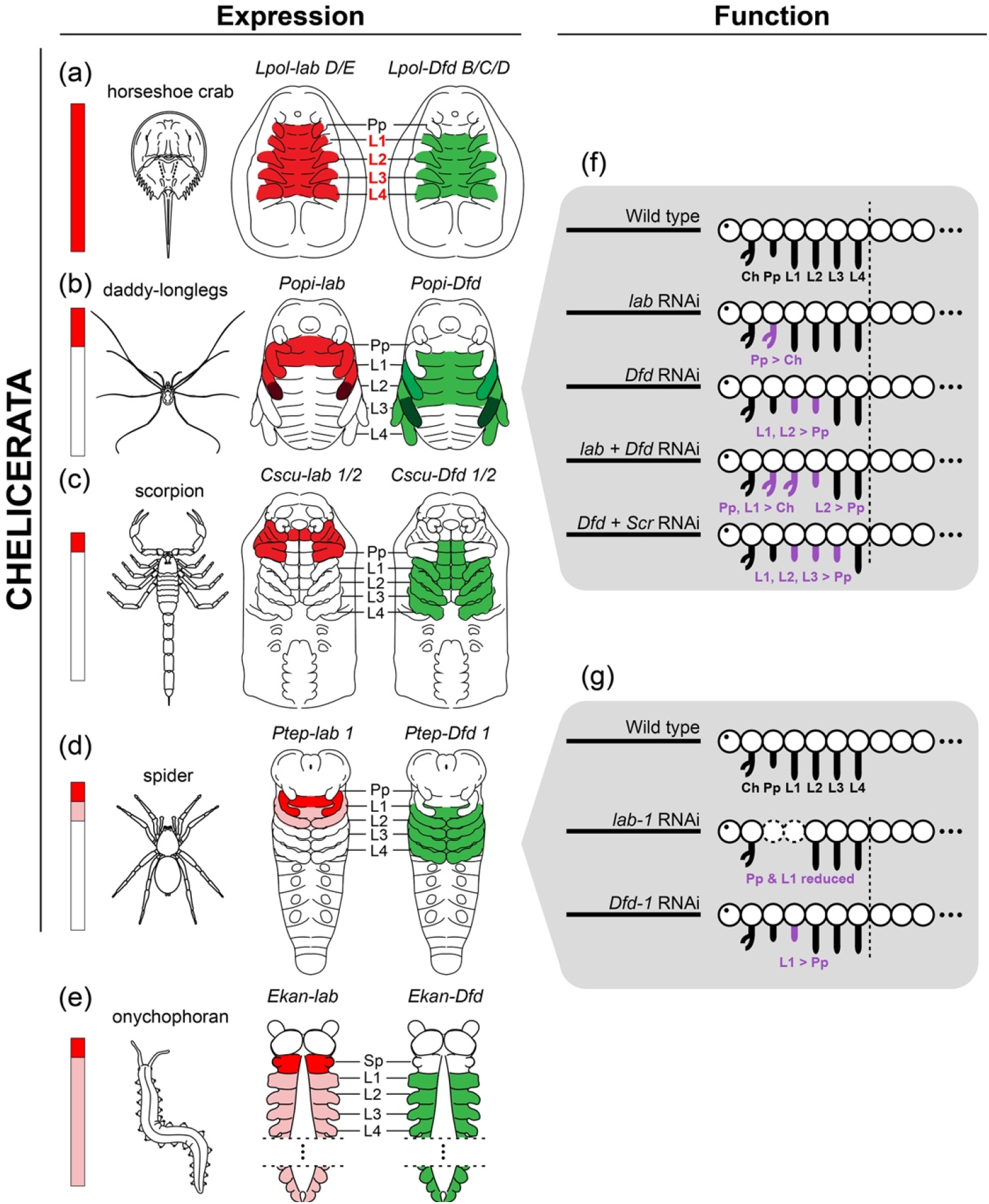
Summary of *labial* (red) and *Deformed* (green) expression and functional data in Chelicerata and Onychophora (outgroup). Darker shades of red and green reflect concentration of expression. Vertical bars on the left portray the degree of *labial* expression concentration on anterior segments. (a) Horseshoe crab *Limulus polyphemus*. (b) Daddy-longlegs *Phalangium opilio*, after Sharma et al. (Sharma et al. 2012a), and Gainett et al. (Gainett et al. 2021). (c) Scorpion *Centruroides sculpturatus*. (d) Spider *Parasteatoda tepidariorum*, after Schwager et al. ((Schwager et al. 2017)). (e) Onychophoran *Euperipatoides kanangrensis*, after Eriksson et al (Eriksson et al. 2010) and Janssen et al. (Janssen et al. 2014). (f) Summary of phenotypic outcomes of Hox RNAi experiments in *P. opilio*. Data for the single knockdown of *Po-Dfd* and *Po-Scr* knockdown after Gainett et al. (Gainett et al. 2021). (g) Summary of phenotypic outcomes of *Ptep-lab-1* and *Ptep-Dfd-1* RNAi experiments in *P. tepidariorum*, after Pechmann et al. (Pechmann et al. 2015).

A refined understanding of anterior Hox function in this chelicerate led us to revisit patterns of morphological disparity across the prosomal segments of Chelicerata more broadly. In the harvestman, each prosomal segment exhibits a distinguishable identity; in the walking legs, this is observed as unique counts of tarsomeres (tarsal articles) associated with each walking leg pair. By contrast, the walking legs of spiders and scorpions are morphologically more similar, being mainly distinguished by allometric differences (e.g., walking legs become longer towards the posterior in scorpions; the first pair of walking legs is the longest in most spiders). Yet, like the harvestman, the pedipalps of both spiders and scorpions are clearly morphologically distinct. Horseshoe crabs exhibit another condition still, with no morphological distinction between the tritocerebral legs (or “pedipalps”) and the posterior three pairs, except in adult males (where the tritocerebral appendage is modified into a clasper for reproduction). Only the posterior-most pusher leg has a distinct, unique morphology by comparison to anterior leg pairs in both sexes of horseshoe crabs.

Our gene expression surveys revealed an intriguing correlation between the presence of a morphologically distinct tritocerebral segment identity (e.g., pedipalp; slime papilla) and the concentration of *labial* expression in this segment. The only arthropod surveyed to date that has strong, uniform *labial* expression on all post-deutocerebral appendages is the horseshoe crab *L. polyphemus*. This condition correlates with the morphological similarity of their tritocerebral appendage and post-tritocerebral prosomal appendages (i.e. these five appendage pairs all resemble pedipalps, or all resemble legs) (Fig. 6a). The only comparable case is the protonymphon larvae of sea spiders (Pycnogonida), where *lab* is uniformly expressed in the third and fourth head segments, which bear anatomically identical larval appendages at this developmental stage (Jager et al. 2006).

In contrast to the horseshoe crab and sea spider larvae, the *lab* homologs of all other arthropods with a morphologically distinct tritocerebral identity (second antenna, pedipalps, or intercalary segment) exhibit anteriorly restricted expression domains, with highest concentration of expression levels in the tritocerebral (and in some cases, also the fourth head) segment (Fig. 6 b– d). This correlation further holds for the onychophoran *Euperipatoides kanangrensis*, in which *lab* is more strongly expressed in the tritocerebral segment and its appendage (and weakly in all posterior segments), correlating with the unique identity of the slime papillae in contrast to posterior lobopods (Fig. 6e) (Eriksson et al. 2010; Janssen et al. 2014).

Furthermore, we discovered that *Dfd* homologs exhibit a similar trend: Morphologically similar legs exhibit uniform expression of *Deformed*. Although *Dfd* homologs always bore an anterior boundary in the fourth head segment, expression patterns and transcript levels of *Dfd* copies were highly similar in the L1–L4 segments of spiders (Schwager et al. 2017) and scorpions (this study); and in L1–L4 of horseshoe crabs. This correlation holds upon inclusion of Hox surveys (albeit fragmentary datasets) from other chelicerate groups; uniform expression levels of *Dfd* were detected in the L1–L3 legs of the hexapod larva of an acariform mite (Telford and Thomas 1998). In contrast to these groups, expression levels of the single-copy ortholog of *Dfd* in the harvestman are markedly heterogeneous across L1–L4, with a unique expression pattern and level of intensity in each leg (Gainett et al. 2021). The general correlation we observe across these groups is that homonomous blocks of segments (i.e., segments bearing morphologically similar appendages) tend to exhibit similar combinations of Hox expression levels.

These patterns suggest that modifications of pedipalp and leg morphology across Chelicerata may be attributable to modulation of Hox levels, driven by variation in cis-regulation of *lab* and *Dfd* to achieve unique combinations of transcript levels in morphologically distinct appendages. The presence of Hox duplications in Arachnopulmonata, while intriguing from a macroevolutionary perspective, may not be the mechanism that underlies morphological disparity of chelicerate prosomal architectures. We found no evidence that prosomal Hox genes have achieved additional segmental identities in arachnopulmonates by spatial subdivision of paralogs’ expression domains (in contrast to opisthosomal Hox copies in spiders and scorpions; (Sharma et al. 2012a; Sharma, Schwager, et al. 2014). This inference is further underscored by the marked morphological disparity between the walking legs in some groups of acariform mites, which are comparably diverse to spiders, but are not part of the shared genome duplication at the root of Arachnopulmonata (Schwager et al. 2017; Ontano et al. 2021).

These hypotheses could be tested further by investigating arachnopulmonates with extreme modifications of specific appendage pairs, such as Amblypygi (whip spiders) and Uropygi (vinegaroons), groups that exhibit marked elongation of the antenniform first walking leg pair. Recently developed tools for the whip spider *Phrynus marginemaculatus* may prove useful in this regard (Gainett and Sharma 2020). With regard to establishment of specific leg segment identities in the harvestman, advanced tools for misexpression of Hox genes are required in this system for nuanced understanding of Hox transcript combinatorics. Ectopic expression of *lab* in posterior segments (L2–L4), for example, is central to testing the hypothesis that co-expression of *lab* and *Dfd* underlies L1 fate specification. Similarly, misexpression of *Scr* in L1 and L2 is required to test whether co-expression of *Dfd* and *Scr* is sufficient for specifying L3 fate.

## Materials and Methods

### Embryo collection and gene identification

Adult *Phalangium opilio* individuals were collected from Bascom Hill, Madison, WI, USA along the exterior walls of nearby buildings. Gravid females of the scorpion *Centruroides sculpturatus* were hand collected from sites in Arizona by citizen-scientist collaborators. Embryos of the horseshoe crab *Limulus polyphemus* were collected in Woodshole-MA, USA in June 2022. Embryonic staging nomenclature follows Gainett et al. (Gainett et al. 2022) and Sekiguchi et al. (Sekiguchi et al. 1982). Details of collection, maintenance and fixation of embryos are described in the Electronic Supplementary Methods.

The complete sequences of *P. opilio* Hox genes *labial* (*lab*), *probocipedia* (*pb*), and *Deformed* (*Dfd*) were previously isolated from the *P. opilio* genome (GCA_019434445.1;(Gainett et al. 2021)) (Table S1). The *engrailed* (*en*) ortholog in *P. opilio* was identified by Sharma et al. (Sharma et al. 2012a) (Table S1). The scorpion *C. sculpturatus, lab* paralogs (*lab-1, lab-2)* were identified from the reference genome (GCF_000671375.1; (Schwager et al. 2017)). The numerical nomenclature for the scorpion and spider (Arachnopulmonata) genes was used to differentiate paralogs from their shared whole genome duplication (WGD) from the independent Hox paralogs of the horseshoe crab WGD. Horseshoe crab Hox paralogs are here referred to with alphabetic nomenclature, following the convention established for *Carcinoscorpius rotundicauda* and *Tachypleus gigas* genome assemblies (Shingate, Ravi, Prasad, Tay, Garg, et al. 2020; Shingate, Ravi, Prasad, Tay, and Venkatesh 2020).The horseshoe crab *L. polyphemus labial* paralogs (*Lpol-lab A, Lpol-labB, Lpol-labD, Lpol-labE*) and *Deformed* paralogs (*Lpol-Dfd A, Lpol-Dfd B, Lpol-Dfd C, Lpol-Dfd D, Lpol-Dfd E*) were retrieved from the reference genome annotation (GCF_000517525.1; (Kenny et al. 2015; Battelle et al. 2016). Accession numbers (Table S2) and phylogenetic analysis of *labial* (Supplementary Fig. S4) and *Deformed* (Supplementary Fig. S5) homologs in the horseshoe crab species are described in the Electronic Supplementary Material.

### RNA interference (RNAi) via double-stranded RNA (dsRNA) embryonic injections

Gene cloning and dsRNA synthesis are detailed in the Electronic Supplementary Methods (Table S3). For the single knockdown of *Po-lab*, two clutches of embryos were injected (n=194). Three clutches were injected in the double knockdown of *Po-lab + Po-Dfd* (n=217). An additional three clutches were injected with *Po-lab* dsRNA, and an additional two were injected with *Po-lab + Po-Dfd* dsRNA which were then fixed and exclusively used in both colorimetric and fluorescent *in situ* hybridization. Additionally, two clutches (n=157) were injected with deionized water as a negative control. Injection mixes were prepared with Rhodamine dextran (1:20) for visualization.

### *Colorimetric and fluorescent* in situ *hybridization*

Colorimetric *in situ* hybridization gene expression assays of *P. opilio* used sense (control) and antisense RNA probes labeled with DIG RNA labeling mix (Roche, Basel, Switzerland) and followed published protocols (Sharma et al. 2012a). Images were obtained on a Nikon SMZ25 fluorescent stereomicroscope with a DS-Fi2 digital color camera utilizing Nikon Elements software. Fluorescent *in situ* hybridization followed a modified version of the Molecular Instruments (Los Angeles, CA, USA) hybridization chain reaction (HCR) v.3 protocol (Choi et al. 2018; Bruce et al. 2021). HCR probes for Hox paralogs were designed avoiding regions of high similarity to circumvent non-specific binding. Probe sequences were designed by Molecular Instruments or in an open-source probe design program (Kuehn et al. 2022). Details of probe design and probe sequences are available in the Electronic Supplementary Methods. Imaging was performed on a Zeiss 710 and Zeiss 780 confocal microscope at the Newcomb Imaging Center, UW-Madison, USA. Z-stacks were projected with maximal intensity mode, and linearly adjusted for brightness and contrast in FIJI (v. 2.9.0/1.53t). Figures were assembled in Adobe Illustrator 2022.

## Supporting information

Electronic Supplementary Material

