## Supplementary material for "Dual functions of *labial* resolve the Hox logic of chelicerate head segments": Electronic Supplementary Material

This PDF file includes:

Supplementary Methods
Supplementary Figures S1 to S5
Supplementary Tables S1 to S3
Supplementary References

### Supplementary Methods

#### Animal Husbandry and fixation

Adult *P. opilio* individuals were collected from Bascom Hill, Madison, WI, USA along the exterior walls of nearby buildings between the hours of 9:00AM and 5:00PM periodically from May to August 2021. The animals were housed in simple rectangular plastic containers in a ratio of one male to eight females. Each container was supplied with wet cotton as a water source, and small pieces of egg cartons to provide shelter. A supply of fresh fish flakes was constantly maintained, occasionally supplemented with frozen crickets *ad libitum* (*Acheta domestica*). Small petri dishes containing moistened coconut fiber were included in the containers to provide substrate for egg laying. These dishes were checked for embryos daily. Egg-laying dishes with embryos were incubated at 26°C for further development.

*P. opilio* embryos used in both colorimetric and fluorescent *in situ* hybridization were dechorionated in commercial bleach for 10–20 min rinsed several times in 1X PBS. Embryos were fixed in equal volumes of heptane and 4% formaldehyde (Sigma Aldrich) in 1x PBS, in the phase between the two solutions, either overnight (colorimetric *in situs*) or for two hours (fluorescent *in situs*). The embryos were then rinsed with PBST (0.1% Tween-20 [Sigma Aldrich] in 1x PBS), before gradual dehydration in methanol or ethanol.

Embryos of the horseshoe crab *Limulus polyphemus* were dechorionated in bleach and fixed for 40min in 3.2 % paraformaldehyde (PFA) in PEM buffer, following dehydration in methanol.

Gravid females of *Centruroides sculpturatus* were anesthetized with Co2 and dissected under 1x PBS. Embryos were fixed for 20 min in 4% formaldehyde solution in 1x PBS, following washes in PBS-T and dehydration in methanol.

#### Orthology Inference and Phylogenetic Analysis

To determine the orthology of *L. polyphemus lab* and *Dfd* Hox homologs with respect to *C. rotundicauda* and *T. gigas*, selected *labial* homologs were compiled from the Homeobox dataset in Ontano et al. (Ontano et al. 2021) and complemented with the *P. opilio labial* sequence from the genome (Gainett et al. 2021), and the *labial* homologs of the horseshoe crabs *Tachypleus gigas* and *Carcinoscorpius rotundicauda* (Shingate, Ravi, Prasad, Tay, Garg, et al. 2020; Shingate, Ravi, Prasad, Tay, and Venkatesh 2020). Accession numbers are provided in Table S1 and the protein alignment is available in the online repository. Four *labial* and five *Deformed* homologs occur in the genome of *L. polyphemus*. However, *Lpol-lab B* has an unusually long sequence, and, despite having a *labial* homeodomain, appears annotated as DNA-excision repair protein ERCC-6-like. It is unclear if this sequence is misassembled or is a pseudogene.

Sequences were aligned as proteins using Clustal Omega (Sievers et al. 2011) implemented in SeaView v. 5.0.4 (Gouy et al. 2010), and analyzed under maximum likelihood with IQTREE v. 1.6.12 (-m TEST -b 1000 -wt; model JTT+F+G4 for *lab*; model JTT+I+G4 for *Dfd*) (Nguyen et al. 2015).

### Cloning and Double-Stranded RNA Microinjection

*Po-lab*, *Po-pb*, and *Po-Dfd* fragments were amplified via PCR from cDNA using gene-specific primers designed with Primer3 v. 4.1.0 and appended with T7 ends (Table S2). Products were cloned using the TOPO® TA Cloning® Kit One Shot® Top 10 *Escherichia coli* following manufacturer protocol (Invitrogen, CA, USA). Clones were Sanger sequenced to verify the identity of the fragments (Table S2). Double-stranded RNA was synthesized using the MEGAScript T7 transcription kit (ThermoFisher, MA, USA) from a plasmid template using T7/T3 RNA polymerase (New England Biolabs, MA, USA) following the manufacturer's protocol.

Double-stranded RNA concentration was verified using Nanodrop ONE spectrophotometer (Thermo Scientific). *Po-lab* dsRNA was adjusted to a final concentration of 4 μg/μL. For the double knockdown of *Po-lab* + *Po-Dfd*, the genes were mixed in appropriate volumes to produce a concentration of 2 μg/μL for each gene, yielding a total concentration of 4 μg/μL. For each microinjection, the dsRNA was mixed with Rhodamine Dextran B dye (Thermo Fisher) at a ratio of 20:1 to visualize the volume of injected solution.

Embryos of *P. opilio* were selected for injection after 4-5 days of uninterrupted development, ensuring proper formation of perivitelline space large enough for needle insertion without contact with the developing embryo. This coincides with embryos at stage 6 and 7 (Gainett et al. 2022). Clutches were first immersed in a 50% bleach solution and routinely agitated to remove the chorion. Suitable embryos were then transferred to plastic coverslips coated with a solution consisting of double-sided tape dissolved in heptane. These coverslips were then taped to the bottom of a small petri dish and embryos were allowed to dehydrate over the course of several hours. Once dehydrated, the slips were submerged in halocarbon oil 700 (Sigma Aldrich). Microinjection needles were produced from glass capillaries using a micropipette puller (Sutter P-1000).

### Phenotypic classification in the RNAi experiments

Phenotypes resulting from dsRNA injection were scored upon hatching, or in the case of severely detrimental phenotypes, upon assisted hatching with manual removal of the vitelline membrane with forceps.

For the *Po-lab* experiment, the phenotypic were classified in (1) homeosis only; (2) homeosis + body segment fusions; (3) wild type; and (4) dead. Homeosis consisted of pedipalp-to-chelicera transformation. Weak homeotic phenotypes showed fusions or defects in the boundaries of the tarsus, tibia, and patella. Stronger homeosis had three segmented appendages, with an ectopic opposing claw. Body segment fusions were scored based on the fusion of the proximal part of the tritocerebral appendage with either the chelicera, or the first pair of legs.

In the *Po-lab* +*Po-Dfd* double knockdown, hatchlings were initially classified into the same phenotypic classes. In this first classification, homeosis was scored as either transformation toward cheliceral identity (as in *Po-lab* single KD), or as transformation toward pedipalp identity (as in *Po-Dfd* single KD (Gainett et al. 2021). Body segment fusions occurred between the tritocerebral appendages and the chelicera or L1 leg.

The 43 hatchlings displaying RNAi phenotypes were further classified in (1) *Po-lab* single KD phenotype; (2) *Po-Dfd* single KD phenotype; (3) *Po-lab* +*Po-Dfd* additive phenotype; (4) *Po-lab* + *Po-Dfd* synergistic phenotype. In the additive phenotype hatchlings showed the combination of the *Po-lab* single KD phenotype (pedipalp-to-chelicera) and *Po-Dfd* single KD phenotype (L1 and L2-to-pedipalp (Gainett et al. 2021). The synergistic phenotype consisted of a transformation of both the pedipalp and L1 leg towards cheliceral identity. We did not observe truncations in L2 besides the reduction of the metatarsus, which we interpret as a transformation towards pedipalp identity (as seen in the *Po-Dfd* single KD (Gainett et al. 2021).

**Hybridization Chain reaction probe design**

*Probes for* Phalangium opilio

*Po-lab* (Table S4) and *Po-en* (Table S5) probes were designed with the HCR Probe Maker tool (Kuehn et al. 2022). Partial sequences were selected based on manual inspection of alignments of available nucleotide sequences with consensus between the genome and the transcriptome sequences available. The following fragments were used for probe design:

```
>Popi-lab_Contig9232_pilon_CDS_labial
ATGAATTCAAATGTAGTGTACGGCGGAGTTTGCCATAACGGCGACGCGTCCTCCGCTTACGGCAACCATCATCATCATC
AATCGCCGTACGGCCACGCCGACCTGGCCGCTCAGGTGTACTACCCTCAACCCGCCGCCGATCCCTTGGCCGGAGCAGC
GGCCGCCGCGGCAGCCGCAGCCGCCAATCATCACTATCTCGGTGGAGGACTAGTCGGAAGCTACGCCACGACCGACCA
CAGGACGCCTCCGCTTCACGGCGACACGGCCAACGGTCAGTTACCTCACCAACACATCATAAACGAACCCAACGGTTT
AAGTTACACTAATCTCGATCAACAAAACGCCTTCGCGCACCAACAACGTTTGTCTCACTTGAACAGCGGTCACCCGGGT
CCCGTTCCGGGAACGACCGGAAGTCCCGGGATGCCGCCTCATTCCCATCACGGTTACGGACCCCCGACGGATATCGGTT
CCAACGGGGGTGGAGGAATGGTGCACAGAAACACGGTGGCCACGGCTTCCTCCGCGCCCTTCGCCGCCTACAGAAACT
TGGACTACGGAGCCCATCACGGGCACCATCACGGACACGACGTCAATAACGTTCATCATAACGGGTCCGCTCACGTGG
TGGATCCCGCGTGTTCCGTGGCGTTAGGCGGCGATGGCAACAGGACGGTTTCGAGCGGGCTAGCGGCCACCAGCGCGC
CTTCCTACCCTAGCTATTTGGACCCTTTGGGCATACCCAGAAGAAACGGTTTCGGTTACCCGGACACCTACGAAGCCAC
GATAAGGGAAGGGTGCCATCCAAACGGGGGTTACCCTAGCCCAACCGCGTTAGGCAGACCCGGACTGTCTCCTCCTCA
AGCCGCGTCCGTACCAACTTATAAATGGATGCAAGTGAAAAGAAATGTTCCAAAACCAGTCAACAAAACAGAATTCGG
TTTCAGCGGCGGAGGAAATATGGTCGGTAGCGGAGGTAACGGCGGTGGCTTGGGCGGAATCGCGGGTACGGGAATGTG
CGGTGGAGCCGGCGGCGGTCTAAACGGATCGAATCTCGGCAACGGTTTAGCCGGTGGTCCCGGTTCGGGTCGGACAAA
CTTTACGACGAAACAACTAACTGAACTCGAAAAAGAATTTCACTTTAACAAGTACCTGACCAGGGCCAGGCGAATCGA
AATCGCCACCGCCTTGCAATTGAACGAGACGCAGGTCAAAATATGGTTTCAAAATCGTCGAATGAAACAAAAGAAACG
TATGAAAGAGGGTTTAATACCGCCGGAACCCATTTCAACGGACAGTTCTTCTCCCCCGTCATCCGCCCAATCTCCCGGT
CAACCAAACTCAGTCGGCGGTAATGGAGCTTTACTAGCCGGTGGTAGTGATAACAGCCAGTGA

>Popi2_engrailed_comp171093_c0_seq2_piece_ordered_for_HCR
TTAAATATTTCTTGTTTTGAAATAAACTTCGGTCTTTTTTTAAATCAATCGGTAGTCTCGAATCCTGATGCGAAGAAACG
CGCCAGTATTTTCGGGTTTTCTCTTCTCTCCGTTAAAAAATCGCGTGTAGGAAAAGCCAGAGTTTTACTTATTACGAGAG
AGAATCTTTCAATCTTACTCGAGGTTCGAGTCTCTCTTCGGGTAAGACCACCGTAGTCGAGGCGGCGGGGAGAAAGATC
GGACCCCCGCTTCGACCCGGTCTCGAAAAGCGGGCGCTATGGCATTAGATACGATGGTAGAGGCCAAGCGCCGAACGT
```

CGATCAGCAGCCCGGCGCGATCCGACGACTTGGACTGTACGTCGTCGTCCGCGGACAGTTCCCGCGTCGACGATTCAAC
GACGCCGCCTCCTCAGCGAAATTCTTCCTTCCCCGGCGCCGTAGAGAACTTACGGGCGCCTCCGGTCATACCAGCCGTC
GTTGGGGCGCCCTTTTCCAGGCACTCGCTCAAGTTTTCCATCGAGAAGATTTTATCGCCGGAATTCGGCAAAACTCCTA
AACACGGGTCGCCCATCGCGGCCGAAGTCAGAAGGACTCATAACGGACTCGTTCATAATCACCTACATCACGCGGCCG
TGCCCCCGCTCTCCAGGACTGTTCCCACGTCTCCGACGGACTTTTTCTCGGCGGTGACCGGTCCGCCAACACCCGCCAG
CAATAACAATCGCGAACACCATCGCAATCATCACCGGCGCGTTCAACAACCCGAAAGTCCCAGTTCGAACTGTAGTTC
CGTTAACTCCTCCGTTAGGCCTCGGAGCGTCGGCGCCTCCACGCCCGAAACCAGCCAGTCGCCCAAATTGCCCTCCTTG
GCGCCCCAAGAAGACGGCGAAACGGGCAAAAAGGGACCCATTTTATGGCCCGCCTGGGTCTATTGCACACGATACTCC
GACAGACCCTCATCTGGCCCAGGTCCCCGGTCCCGACGCATGAAGAAGAAGGAAAAGAAACCTGATGAGAAGAGACC
CAGGACAGCGTTCACCGCCGAGCAGTTGGCCCGGCTAAAACAAGAGTTCCAGGAAAACAGATACTTGACGGAAAAAA
GACGACAAGATTTGGCCAAAGACTTGAAACTGAACGAGTCCCAAATTAAAATTTGGTTTCAGAACAAACGGGCCAAAA
TCAAGAAGGCCAGTGGACAGAGGAACCCATTGGCTCTACAGTTAATGGCCCAGGGCTTGTACAATCACAGTACCGTCG
CCATGGATGGCGAAGGTGAAGATGATTCAATGGACAAATCATAG

*Probes for* Centruroides sculpturatus

*Cscu-lab 1* is shorter than the other paralog, and only part of the UTR sequences could be verified, because the transcriptome sequence is also fragmentary. A fragment of the sequence was provided to Molecular instruments that includes CDS and part of 3’UTR that has a perfect overlap between the genome sequence and the transcriptome. It also has part of 5’UTR. Nine probes pairs were designed, avoiding a repetitive portion of the UTR.

>Cscu_lab-1_CDS_some_3UTR_XM_023365857.1 PREDICTED: Centruroides sculpturatus homeobox protein Hox-A1-like (LOC111623321), mRNA
TAATGTGTAAAGCGCCAAAGTAATTTTAGCGATAGAGAAGTGTTTTTTTTTTCGATTTTTATTGAAGTTTTACCTAAAAA
CACCATCAAACTAAAACCTAAATGTAGAATTTATAGGAAACTAAACTTAAGTCTCGATATATTTGATTTTCTTTCTAATG
TTTCACTAATAGCTACCATTGTTGACAATGTATAATGGGTTCTCTTCTATGTCTCTCTCTCTCTCTCTCCTTCTCTCTCTCT
CTCACTCGTTCTGCCTGTCTGTCTGTACCATCAGTCAACAAGAGTGACTACGGCTTCACTGGTGGAGGTAACCAGGCTT
GTGCAGGTGCAGGGCCAGGATCGGGGCGTACGAATTTCACGACCAAACAGTTAACAGAACTTGAGAAAGAGTTTCACT
TCAACAAATATTTGACGAGGGCAAGAAGGATCGAAATTGCCACCGCCCTTCAACTTAACGAGACTCAAGTGAAAATCT
GGTTCCAGAACAGGAGAATGAAGCAGAAGAAGAGGATGAAAGAGGGCCTGATACCCGGTGAACCCGCCTCGAAGGAA
AACAACAGCCCCACCTCATCCTCAAGCCCTCCTAACAGTGGCTCCGCCCAAAATCAGACTGCACTTACTTCAAAGGATG
CTAACCAGTGAGCGAAAGTAATGTGTGTGCGTGTGCGTGTGTTTTTTCGCGGCTGTTTTCGTCGTACTCCGAACCGAATT
ATGTACAGATGATGCCGAAGATGATGAAGATT

*Cscu-lab 2* is longer than *Cscu-lab 1*. A fragment of the 5UTR and the CDS was included in the design and validated with the genome and transcriptome. This sequenced was provided to Molecular Instruments, and 15 probe pairs were designed.

>Cscu_lab-2_some_5UTR_CDS_XM_023366700.1:132-1250 PREDICTED: Centruroides sculpturatus homeobox protein Hox-D1-like (LOC111623958), mRNA
TTTGCTGATTTATCTGCTGGGGGAGCGAGACGGGCAATTTCCTACCGTCTCGCTAAAAACTTTACTCAATAGATACAGA
CTGGGGGACATGAATTCTAATGTCGTATATCCTTCCATGTGTAGAAACGGCGATGCTGGAGCGACACCGGGCTACGGG
ACTCAGGCATACGGAGGCAGTGGCGGAAGCTCTACCGGGTCGACGGGAGAACTCTCCGCTCAGGTGTATTACCAACAA
GCGGATCCGCTGACAGCAGCAGCTAACCACTATGGCTTGGTCGGCGGATTCGGGACCCCGGATCCACGGACTCCACCG
CTTCATGGTGAAGCCACTAATGGACACGTCCCGCCGCATCATCAAATCATCAGTGACAATAACGGACTAAGCTACACT
AATCTCGATCAGCCGGGATATGCCGGACAGCAACGGGTGCCACATCCTCACCATCCCGGTTACGGTACAACCCCTGCA
GATGCCCGGCACCACCACGGGTTTGGCTATCGTCAGTTGGAATATCCTCACGACGTGAATGCGTTACACCATCCAGCTG
ACCATCAGGGACCTGTTTTGAGCGCTGATGGTGGAAGACTCGTTACAAGTGCCCTACCTCCACCCTCTAACACTACTTA
TCCGTATTTAGATTCATTGACACTGACTAGGAGAAACGGATACGGCTATGCGGACGGCTACGATGCGGCTATGCGAGA
AGGATGCCAGGCAAACGGCGGCTACAGGACGTCTGCAGCCATGGCTAGGGCCAGGCTATCTGCGCCTCAGCCGACACC
GGTACCTACCTACAAGTGGATGCAGGTGAAAAGGAGTGTGCCCAAAACAGTGAATAAGACCGATTACGGTTTTAGTGG
TGGAGGTACGGGCACCAATGGGGCAGCTGCAGGTCCAGGTGGAACAGGTAGGACCAACTTTACAACAAAACAACTAA
CAGAACTCGAAAAAGAATTTCATTTCAACAAATATTTAACAAGAGCAAGAAGGATCGAAATAGCCACAGCTCTTCAAC
TAAATGAGACCCAAGTCAAGATATGGTTCCAAAATAGAAGGATGAAGCAAAAGAAAAGAATGAAAGAGGGTTTAGTA
CCGCCGGAAAATCCGACGGGTGACGTTACAACATCGAATTCGGGTCAACAAAATCCAACAGTTTCACCTAATTCGGGA
GGAACAGCGACGACGCAAAAAGATTGCAGCCAATGA

A partial CDS of *Cscu-Dfd 1* with partial 3'UTR was used for the design. This fragment was provided to Molecular Instruments, and 14 probe pairs were designed.

>Cscu_Dfd-1_CDS_some_3UTR_XM_023374825.1:1-1107 PREDICTED: Centruroides sculpturatus homeobox protein Hox-D4a-like (LOC111630686), mRNA
ATGACGTTGATAGAGTTAGAAGAGGAGGATATGTTGGAAAAAAAGGATGTTGTTAGAGTTAGGAAAGATGCTAAAGAT
GAAGAAAACGAAGAGGAAACAGCCATTTTGCCGTCTACCCTGTCCAGGCGAGCGAGAGCTCATCGATCATCCGTGACG
TGGGTCAGCGCCGCTTTACCAGATTGGACGGCAGCCGTCACGTGTTATGCTAATTTTTTGCCCGGGGGCGGAGTCGCGG
AGCGAGTTAAACGACAACCCGAAAGAAAAATTATTTCCATCCAATTCACCCAAAAATTAATGATCATGAGTTCGTTTTT
GATGAACTCTCCTGCGTATGCCGACCCGAAATTTCCACCCAGCGAGGAGTATTCACAGGGCAACTACATCCCCAACCAC
GCAGCCGACTACTACGCGCCCCCGCCACCTCAATCGCATCCTTACAGTTACGGCTGTGTCCCTCATGCAACCTCCACGC
AGCAGACTTTCGGCCATGAGGCATCACAATACAACCACCACGCCGGTCCCCAATTCCATCAGCAATATCATCAGCAAC
CGTGCGGTTTGGCGGCGCCCCAGGACGGCAACAGGAGCCAAAACTTGGCACGGGGGCCCCAACTACACCACCAAGCTC
AACAGGAAGCGGAATGTAATACAAGTCAACCCCTCATCTACCCGTGGATGAAGAAAGTACACGTCAATTCAGCTAATG
GGAACTTTTCGGGTGTGGAACCAAAAAGACAGAGGACCGCTTATACTAGGCATCAGATACTCGAACTGGAGAAAGAAT
TCCATTTCAATCGCTACCTGACGCGTAGGAGAAGGATCGAAATTGCTCACTCCCTTTGCCTGAGCGAACGACAGATTAA
GATCTGGTTCCAAAACAGGAGGATGAAATGGAAGAAGGACAACAAATTACCCAATACTAAGAACGTTAAGAAGAAGA
ATCCCGATGGTACCACGTCCAAGAGGTCTCAGAATTCTAATCCGGCAGGGATGGTAACACATGCGGATACTGTGCCAC
CTCCGCCACCCATGCACATGATGCAACCTCAGCTACCCCCTCCACCATCTATGGACTCCAAAAGTGGTTATGGGCTGAC
AGAATTATAAAATGGATAAAAACCCCTTCAATACAGAGAAAGAAAGTGTAGAGAAGTCGTCGATGTGTGCTAGTTTAT
AGTACTTCGCCTCTACTTTTTGGTGCTTTAACAAA

A partial CDS with partial 3UTR was used for *Cscu-Dfd 2* paralog. This fragment was provided to Molecular Instruments, and 18 probe pairs were designed.

>Cscu_Dfd-2_partial_CDS_some_3UTR_XM_023379077.1 PREDICTED: Centruroides sculpturatus homeobox protein Hox-B4-like (LOC111634317), transcript variant X1, mRNA
AAATTAATGATCATGAGTTCGTTTTTGATGAACTCTCCGTCTTGTGTGGATCCTAAGTTTCCACCAAGTGAGGAGTATTC
GCAAGGTAACTATATACCGACACACGGTGGTGACTATTACGCTCCCCAACCGCATCCTTACGTCTATGCGGTCAACCCA
CCGCCGCCTCATCCCGCTGTTACGTACGGACCTGTGGAAAACGGTGGAGGTCCGTGTTATCCCGATCACGGCGGCTCTC
CCCTCTACTACCAGCATTCCTGCAGCATGACACCGAGTATCGGAAGCCGGGTCGCAAGACCGACACCGGACCACGTTA
CGTCCAGCCACAGTCCCGTTACGATCCTTCCTCCCGGTCCAACTCGCAGATCGCCGGTATCTTCGCCTCCACCGGGCGTC
GTTGGGCTCAGCCGGGTCCAGCAATCGCAGCAGCCACAGCAGAATCAAGCCCAACAACCACCTCAACTTAGTCAACAG
CCGTCACAACTCCATCAAAGTGCAGCGAGTCAGCCCGATACGGGAAATCCCGACTGTGCCGTTTCTGGCGGTGGCCAG
CCCGTCATCTATCCCTGGATGAAGAAAGCCCACGTGGGTACAGCTGCAAATGGAAACTTTACTGGAATGGAACCTAAA
AGACAGAGGACAGCGTACACTAGACATCAAATTTTAGAACTTGAAAAGGAATTCCACTTCAACCGTTACCTGACGCGG
AGGAGAAGAATCGAAATAGCTCATTCCCTGTGTCTCAGTGAAAGGCAGATAAAAATATGGTTTCAAAATAGGAGAATG
AAATGGAAAAAAGACAACAAATTACCGAACACGAAAAATGTTAAAAAGAATCCAAATAGAAATCAGAATTTAGCTGC
TATCAATAATCGTAATCAACAGAATCAACAGCGACATCAAGCACCACCACAGGCCCAACAGCAACCTCCATCTACATT
GCCGCCACCTCATCATCCACCAACACATGATCCAAAAGACGACTATGGTCTTACAGAGCTATGATGTGTATTACTCAAC
ATTCCAACTCAGTTACAAGTTCTCAGAATGCGAAGTATATGGGACATGGGAGGGGGCATGAGTACCCTTCTCAACTCTC
CCACAAAACTCCCGCCCCCTGCAATATATCTTCGCAGTGTACTGTACTCAGGAGCAAAGTGCAGAAGCTGAATAGTTTG
TTGCTGTGCATGTGACGCCGTCAGGAGATTGTAGTCGCTAGAATATAACGCTAAGGGATTTTACACGATGAATACCAGG
CGGAACCACAGGACACAGTCGTAGCAAAGGCTCAGACTTTGCTGACTTAACTTTGGCTAAAAGATCTAAATTTCGAAA
ATAATATTGATTTCAAATTGTTCTTTTGTAAAAAATCTACGAGAAAATTGTTTATTTAAAAGAGGTTTGTAATGTGGAGA
AAATGCAAAGCGGTCAGCATAGAAGAACCTAAGATTCTACCCTTGTCGTCCAATAACTGCTCCTG

*Probes for* Limulus polyphemus

Probes for *L. polyphemus labial* paralogs (Table S6, S7) were designed with HCR Probe Maker tool (Kuehn et al. 2022). Given the high similarity of the homeodomain nucleotide sequence and adjacent sites, this portion was excluded from the design to avoid crosstalk. We did not design probes for *Lpol-lab A* (short sequence) and *Lpol-lab B* (unusual sequence; see "Orthology Inference and Phylogenetic Analysis").
The following sequences were used for the probe design of *Lpol-lab E* and *Lpol-lab D*:

*Lpol-lab E*

>Lpol_labE_XM_013934892.2 LOC106474199 [organism=Limulus polyphemus] [GeneID=106474199]

TAGGAGATATGAATTCTAATGTTTTGCATTCCAGTGCCTGTCACAACAACCAGACTGGAGCATACGATCCGCCCTTTTA
TGGACCTGAATTAACGGCTTCCCATGTTTTCTATGGACAGTCCTCGGGACATCTCTCTCTACCAAATAACCACTATGTAG
GGAACGGTCTGGTCACTTCTTTTCCTGGACTTGACCATCGGTCTTCCTTGCTTAATCGAGAGGGAAGTAGTCAGTTGTCA
GTGCATCTCGCAAGTGATACATCCAGTGGAATGACCTACACAAACCTAGAAAACAACGTGAACTATGTCCACCCGAGT
CGGTTAAGACACTCTCAAGTGTCCAACAGCGGAAATCAGACCATCGCTAGAGTAGAATATGCCGCCACATCGACAGAC
GCCGATAGGGGAAATAGAACTCAGCTTGCCGAAAATTTCACTCTCAGGACCATGGAGTACGGACCAGAAGCAAAAGGT
TCTCGTGGTACGACCCACCATCACCAAACCCTCGTTCTCGAGAGTACAGATGGTCGACGCGCCGAGCTCGATGGTCTCG
GCATCGGAACTCCCGTGAACTCCCATCCATGCATGGCTCCAATGGGGCTCCCGCGGAGGAATGGATCCGGCTACCCGC
AGGGCTATGACGCCCATCATCGGGACACCTGCCAAAGTGACAGCGGTATTTCAGAAGAAGCTGTTATCACCACCAAAA
CTTCGACTGTTCAGCACTCTGTTCCCGTCCCCAAATATAAGTGGATGCAATTTAAGAGAAATATGTCAAAAAGTGGCCA
CAAGTCGGTCCACGAATACAACAGGCTAGGACCTGGTAGTTGCGGCCCTGGGGGAGAACAGTCAACGACCAATGGGCC
AGGAAGAACAAATTTTACTACCAAACAGCTAACGGAATTGGAGAAAGAGTTTCACTTCAACAAATACCTGACCCGAGC
GCGTCGTATAGAAATTGCAAACGTTCTTCAATTGAACGAAACTCAAGTAAAAATTTGGTTTCAAAACCGACGAATGAA
ACAGAAGAAAAGAATGAAAGAAGGTCTCATTCCATCGGAAACTGTTAACCCAGAAATAACTTCTACGTGCCACGAGCG
AAAAATGGTAACTAGGATAACTCTGTCAGGCGTTTCCGCAGCAACAGTATTACCAGCAACTAAGGGAACCTGCTAGCG
ATAGCGAAGCGGACATTTGTAATTATAACAATAAAAGGTATAGACAGCCAACAGCAAATTATGGCACATACTAACGAG
TTTCATAATATTATAAACAACATTTCCATGGCAACAGTGTTACCAGCGACTTAGGGAATATGTTAGTGAGTACTAATAA
TACTACAAAACACAGATACAGCAACAATACTACTAGAGAATTGTGGAACCTGCCAGCTAGTATCCGTAATGTAGCAAT
CTAAATATACACAGTAACAATGCTACCGAAATTTAATGAACCAGCTAGCAAGTTATCGTAAAAATAGAACAAAACCTA
ATCACAATAACAGTGCTAAGAGCGATTTAAGCATCTGATAGCCAAGTTAAAAAATAAACTGCTGGCAAATTTCATTAA
ATTACAAAATTAGAGGAATTAATTCAGTACCTAGCCGTTAAAGAAAGAGAATTTGGAAGCTTGAGAACATTGTTTTGAT
GAAACAGTGATCTCTTGTTAAGACTATGTGGACCAAGCGGAAAAAAATTGCTACCGACAATTTAAATTTATTATGTATT
TATATATGAATATGTATGTATATGTGTGTACGTGTTTTTGTATGATAAAACCAAATTTCATTTGGTATGTGCTGAACTTG
AATCTACAAAGAATATAATATTAGTAATAATTTTAAGAATTACTCTAATTTTAATTTTTTTTTCTTCCCTGCACACTCATA
CCTAACATAAATATTTCAATTAGAGTCGTTCTCTTTGTCAA

*Lpol-lab-D*

>Lpol_labD_XM_013920641.2 LOC106460883 [organism=Limulus polyphemus] [GeneID=106460883]
AAAGGCGGTTCCGAGGTCATAGTAAAGCGCTATTGGTTGGTAGAATCACGTGGCTTGTTCAACCTATAATTCTGTCATT
TACCGTTCATGATTTACCGTCCATTTATTTTCTGATTTATCTCCAACAACGACAGCTGATTTGTGGCACATCTTCAAAGC
TTTTTTATCTGGAAAGACACTCTAAGAAGTTATGAATTCCAATGTTGTGCATTCCACATCGTGTCATAACAGTCAGCCGG
GAACCTACGGAGCTTCTCTTTATGGACCCGAATTAGCGGCTCCCCACATTTTTTACGGGCAGTCCCCGGAGCATCTCCC
ACGACCAAACAATCACTATGTCGGGAACAGCGTGGTCACTTCTTTTCCTGGTCTTGACCACCGTTCATCCCTGCTTAATC
GAGAGGGGAATAGCCAATTACCAGCACACCTTGCAAACGATACATCTAGTGGAATGGCCTACAATAACCTGGACGGCA
GCGTGCGCTATGTCCACCCAGGGCGTCTCGGACATTCGGAAATTGGGAACAGTGGAAATCACACTCTCAGTAGTATGG
GGTACGTTTCCTCGTCGACGGACACCGGACGATCTAACGCTACTCAAATTGCTGAAAATTTTGCTTTCGGAGTCTTGGA
GTACGGTTCAGAAGCAAAATGTCCTCGTAATGTATCCCCTCATCATCAAAACATGATTATAACGGGTTCAGAGAGTCGT
CGTAACGGGCTTGATGCTCCCGGCGTGGATACTACCACGAGCCCTCATCCTTGTATGGCTACAGTGGGCCTCCCGCGGA
GGAATGGCTTCGGCTACCAGCAGGGATATGAAGCTTCTCACCGGGACACCTGCCAAAGTGACAGCGGCACCTCAGAAG
GTGTAGTTATCACCAGTAAAACCTCGTTGATGGCTCAGCACTCTGCCCCTGTCCCTAAGTATAAATGGATGCAATTCAA
AAGAAACATGTCAAAAAACGCTCACAAGTCAGATCACGAGTACAATGGGATAGGACATGGCGGTTGTGGTGTTGGAGG
AGATTCGTCAGCAAGTAATGGACCAGGCAGAACAAACTTCACTACCAAACAACTAACAGAATTAGAGAAAGAATTTCA
CTTTAACAAATATCTGACTAGGGCACGACGAATAGAAATTGCCAACGCTCTTCAACTGAATGAAACACAAGTAAAAAT
TTGGTTTCAAAATCGAAGAATGAAACAGAAGAAAAGAATGAAGGAAGGTCTCGTGCCTCCAGAAACCATCACTCCAGA
AATTAATTCCACTTGCCATGAACATAAACCCGTTACCAGGATACCTCCGTCGAATGTTTCCACGGCAACGGTTCTGCCA
GCCAATAAAGGAACTTGTTAGTGAACATTGGAACATTGCAAGAAAATAAAAATAAATCAACAATTTCTTTACCGAGCC
GTGAAAAACTACGATTTTTGGATGGATGCTTAAGAACATTTTTGGTAAAACATTCATTTCTTGTAGGACTACTGTGGAC
CAAAACATCAAATTGTATGGGAAAATGAAAAAAACAAACAAAAATGTGTATATATATATGCACGGAAAAATAGATTTT
ATGGGGACCTACATAAGAAGAAAATAACGGTAACTAAATCCTACTTTCATTATTCTGATTTTTCTCACGTATTACTCACA
CGTAAAATTATTAATATGGCACAAAACTGCTCTTGTTAACAAAAACATAAACGTTTTTTCGGTTAAGATTAGGCTAAAA
TAAGTTTAGAAAAGCCGAATTTTAGCATATTCATTCTTTTAGCTGAATAATGAATTCTATATGGCATTTTCTTTATTATTT
CTATAGAGTTGTTTTGGCTTGGCTTAATTTCTTGT

Probes for *L. polyphemus Deformed* paralogs (Table S8, S9, S10, S11, S12) were designed with HCR Probe Maker tool (Kuehn et al. 2022). Given the high similarity of the homeodomain nucleotide sequence and adjacent sites, this portion was excluded from the design to avoid crosstalk. The only paralog for which we did not exclude the homeodomain for the design was *Lpol-Dfd B*, so we cannot rule out that there exists crosstalk with the other paralogs. For *Lpol-Dfd B*, the whole CDS was included in the design. The UTR was excluded because it generated some probes with crosstalk even withing the CDS.

For *Lpol-Dfd A*, *Lpol-Dfd C*, *Lpol-Dfd D*, and *Lpol-Dfd E*, the first 110bp of the CDS was discarded because of high similarity. The design in these cases included UTR to reach 20 probe pairs when possible.

*Lpol-Dfd A*:

>Lpol_DfdA_XM_013928948.2 LOC106468515 [organism=Limulus polyphemus] [GeneID=106468515] [transcript=X1]
TAGAAAACGGTACTGGCTGCTATTGATCACGTGATATGCTAATTTATGGACGTAGGCGGAGTCAAGTCACAGTTAAACG
ACAACCCCAAAGAAAAATAATTTCCATCCAATTCACCCAACAATTAATGATCATGAGTTCGTTTTTGATGAACTCTCCT
CCATATGTGGATCCGAAATTTCCACCAAGTGAAGAATACTATCAAGCAAACTACATTCCAAGTCAGCGAGGGGACTAT
TACAATCCGCCAAGTCATCCCTATCGCTATAGCGGCGTCAACAGTCAGCACGCGCCAGTGAGTTATGGCCACGAGCAC
AGCGGTGCCAACGCAGCGTACACAAATCATGGAAGCCCTCCCCGTTACTACCAGCCGCCGTGTTCTCTACCACAAAACT
CACTCCACAGACCAACAATTTTGAGCCCTTCAGTTGACCATGTACCACTAACCAATAGTCCGAACCATATTTTACAGAG
CCCACCGCAGCACCATCAGAGATCTCCGGTCTCTTCGCCGGTGCCGGCGCCAGTGCCGCCTGCAGTGTCTTGCAACACC
CCACAGCCAACTTCCGATCTACAAGCTCGCGTACCACACCAACAACATATTTCTCCTCAGCAGCAAAGTTTAATAGAAA
CGACTCCGGACTGTGCTGTGTCAACGACGGGACATCCGGTCATCTATCCTTGGATGAAAAAAGTCCACATCAATGCAGT
GGGAACCAATGGAAATTTCTCTCCCGGGGTTGAAACCAAAAGACAGCGGACGGCTTACACGAGACACCAGATTTTGGA
ATTGGAAAAAGAATTTCATTTTAACCGGTATCTTACAAGGCGTCGACGGATAGAAATAGCTCATTCGTTATGTTTGTCA
GAGAGACAGATAAAAATCTGGTTCCAAAACCGTCGGATGAAGTGGAAGAAAGACAACAAGCTTCCTAATACGAAGAA
TGTGAAGAAAAGGAACCAAAACGTAGAACACCTGAACAACCATACGCAACCATTAGCTCCTGTATCAAACCACACGCA
ACCAGAAGCGAGACTTCTTCCCCCAGCGCCAAGTCAGTCAATTTCAATGGACCCTAAACCTGATTATGGTTTGACCGAA
CTCTGATATTTTCTTATGTGTGAAGATCATTCATAAGCTGTGTAGTTCTTGACTAAAATTTGGGCAGCATTTTATTATGTT
AAATGTTGTTCAAGGCCCAGACTGTGGTGTATGTGTGTGGATGTGGTGTGCTTAAGCCTCGCGGTGATTTATGTATATGT
GCAAAAATGATGATGAGAAGCGTTTTTTATTTACAATGATCTCTTAAGCGTCGTTAAGCTTTTTGGTGAGACTAACAGT
CTTTCATATAACAAAATGTGTCTCACAAAAGGAAATGTATTTAGACAGTTTATATGCCTGCTTGTACCCAGCTAGCTGTT
CATTGAACTTGTACAGAAATCTGTGGTTTCAAGGAGAAATCCCTAACAGGTAACAGTTATATGGGAAGCCGGGGTATT
ATTAATGTAACATCTGTAATTTTTTTTTCCATTTTGTGTGTGATACTCAATCTAATTTACAGCTTTTTAACTGGTGTGTAA
TCACTGTTAGGTTGTACCATTCTTTATCGTCTTTATTATTAGTAGGTGTAAATATCTATATAATTATATATAAATATATAT
ATATATATACATATATAAAATTGTGGTCAACATTTTGGCTTCCAGGTCAACAACATGGTCTCAAAACTGGATGTACACG
CAAAAATATTGTAATATGTGCATTTTTGTTTAATTGTGGTGGTTTACGATCTTGAGACAAACTGGCCTGTCTGTGTCTCT
CAGGGTATTTCAGTTTGACAAATTCGTAATTTTAAAGTGATATTTATTCCGATTCTGAGAAAACAAACACTCATTTAATT
ATAATATATCCGTTGTTGTGACAACGTTCTTTGGGTTTGACGGTTGGTACTCATGTAATGAAAATGGTGTTTGTGATGTA
TGTATACTGTTCGTGTGCCATAACTATATGCAGATGATATATCTTTATTTTTATTTCATCGACCGTTTTCGCCGGTTGTAC
GATAACAACAACATTTTTTTCTAAGTTGTCTTAAGACATAGAAAGTTCATTAACTGTGACATTGGTCATTAGCAAATAA
CTGACTAAATTATTTTGTTACGTAAAATATTGATTTTAATTGTAGTCTGACTGCAATTCTCTAGTGTTAAAGAACGAAAG
GTCAACGCTAGAGTACGCAAAGCATAGTCAATTTCATGTATTACTTAATGTTAAATGTACTTGGTGTCAGTGTATTTTCA
GTAATTCTCTGATGTATAGCCCATTATTTTAGTAAAGGGAGTTAAAATGTCGACACGAGACTTCTTTAGTCTCCACGTGC
TACTTAAAAGTCTGGGAATAGGTAACACCGTTGGTGATTTTTAAGTATGTTGTAAGAAACAAGACTGAAATCGGTTGGT
CAGTACCTAGTGGTCATAGATTTGCATTCGAATCTGAAAGTTCTCCTCTTGGCTCTTGTCGTAACATGAGTTGTAATGTG
GTCAGTATTCCTAATTGTTGGTAAAACCTTGTTTGCAGGAAACGCTAAAGAAAAATAATATCGTAATAGAACTCATGAA
AATGATACTGCTATTATGTACCATCGTGCATTCCAAGTAGCAGTAGCGACAAACATGCAACTGTAACCAGATCTATTTT
TGTAGTATTATACTGTTTGCTTCAAGGTCATGACAATGAAACCTGTGTACTTTCTGTTATAAACTATTGTAAACACAATC
TGTATATCATTGCTGTCATTACTATAATAACGTAGGATGCAGTGTCCTTTGTAGCAAATTTATTAGGAACTATGATACAA
TATTGTAGTTTTCCGTTAGATCTCAGAGTGGTGTTTGACATTGAGTTTATATATCTTAATTTCTTCTGTAATTAAAATAAA
TTTAAACAAATCAATTGCCCTAGATTACAGCTAAGAAAGTCTCTTGTTCCTGTATTTTATGTACTTTAAAAAAAAACTTC
AGTCCTTCTACTGCTCTATTGATGTAGTCCAAACAATAACTGTCTTTGCTTCCCTCTGGCTGTACCTCAGGAGAAAAACA
ACAACAACAAAAAACAGGCCAATAAATGTGCTTTAAATTA

*Lpol-Dfd B*:

>Lpol_DfdB_XM_013921271.2 LOC106461448 [organism=Limulus polyphemus] [GeneID=106461448] CDS
ATGACCATGAGCTCGTTTTTGATGAACTCTTCGCCGTACGTGGAGCCAAAATTTCCTCCAAATGAAGAATATTACCAGA
CAAACTACATCTCAAGTGATCGAGGAGACTACTACAGTCATCCGTATCGTTATAGCGGCGTCAACAGTCAGCACGCGTC
AGTGAGTTATGGTCACGAGTACAGCAGTCATAACACGGCTTACACAAATCATGGGAGCGTTCCCCACTACTACCCACC
GCCGTGTTCTCTGCCGCAGAACTCGGTAAGCTCGCTTCACAGGCCAAAGAATTTGATCTCTTCAACCGGCAATGTTCCA
CCTACCATTAGTCCGAACCAGGTCTTGCAAGGCCCAGCGAGGCAGTCCCAAAGATCGCCGGTCTCTTCGCCGTTACCGC
CGCCAGCGCTGACAACCGATACTGACCTACAAATTCACGTATCGCATCAGCAAGAAATTTCTCCCCAAGAAAAAAGTT
TAGTTGAACCAATTCCCGACTGTGCTGTGTCAACGGCAGTGCACCCCGTTATCTACCCCTGGATGAAAAAAGTCCACAT
TAATACAGCAGTAGCCAATGGAAAGTTCTCTCCCGGCTTGGAAACCAAAAGGCAGAGGACAGCCTACACCAGACACCA
GATTTTGGAATTGGAGAAAGAATTTCATTTCAATCGGTATCTTACAAGGCGGCGACGGATAGAAATAGCTCATTCATTA
TGTTTGTCAGAGAGACAGATAAAAATCTGGTTCCAAAACCGCCGGATGAAGTGGAAGAAGGACAACAAGCTTCCTAAC
ACGAAGAACGTGAAGAAAAGGAACCAGAACGTAGAATACCTGAGTACGGCGCCAAGTCACTCGCTTCCAATAGACTCT
AAACCTGGTTACGGAACGTGA

*Lpol-Dfd C*:

>Lpol_DfdC_ XM_013936704.2 LOC106476035 [organism=Limulus polyphemus] [GeneID=106476035]
AGCTTGTAAAAATAGCACTGTTGTACACAAACAGAGTTATCTTTACGTAAAGCTGTTTGATAAACTTGAACGAGAGTGG
ATAAGACAGTTTCTGTAACTAGGTTTATGTTCTTAGTAACCGTAACAAGATTGTAAGATCAAGACTGGAGTTTCTACTA
ATATACAAACACATAAATGGTATACGCTAGACTGAGTATAACATAAGTTTCATAAATAACCTCCTTAGATAATCGACCT
TTCCTACGCAAATAAAAGATCGCAGTGATCTTATTGGTCGGATCCACTTACATGGATAGCTTGTGTTCTCCCATCAAATC
GCATAAAACACGCAGAGTTTGCTGTCTGGAGAGGTATCTTTAACCTGATGGACTTTGATTCCACATTTAAGACTTATAC
TTTACTAAATACTTACAGTCAAACACAAGATTTCCAATAGAACAAATCCAAGATGAACTGAATGCACGAGCTTTTCGGT
AAACGATGCCCAGCGGAATAATTAAGCGACCATTCAACGCTGATTTTATAGGCTAATTGGCATTCGTCTTAACGACCAG
TCTGTTTCTATCCCCCAGACACGTGGCTCACGGTTCTAAACAATAACTCTACAACATTTTTTTTCCATTCATTAAGGAAC
ATATTTTAACAGTTGTTACTCCATACTAAATCTGGTGGAATTATAAAAGTAAACCAGTGCATTCTCGCAGGTAGTCCGC
ACTCGCCATGTTGCACTAATCTATCTCTGTCAACTAGCATTGTGACGTAAGCACGTGTGTCAACGTTAGACGGCGTCATT
GGCTGCTACAGGTCATGTGATATGTTAATTTCTGGAGGAAGGAGGAGTCAAAGCACAGTTAAACGACAACTCCGAAGA
AAAATTATTTCCATCCAATTCCCCCAAAAATTAATGATCATGAGTTCGTTTTTGATGAACTCTCCTCCGTACGCGGAGCC
GAAATTTCCACCAAGTGAAGAATATTTACAAACAAGCTACATGGAAAGTAACTACTACAACCATCCGTATTGTTTTAAC
GGCCAGCACACGTCTGTGAATTATGGCCTCGACAACAGCGGTGGCTACACAAATCATCGAAACTCTCCTCATTACTACC
GGCCTCCCTGTTCTCTCGCGCAGAACGTTCACGGGCCAACGAGTGGGAGTTCCATAGTTAACCCGACTCCGGCTACCGA
CAGCCCGAACCGTGCCTTGCAAGGGCAGCCGCAACACCCTTATAGATCGCCGGCGTCTCCGCCAGTAGCGGCCCGTAC
CTCGTCACCTAATCAGCAGCAAGGGTTTACCGAAAAAAGTCCCGACTGTAACGCGTCGACGACGGGACAGCCGGTTAT
CTACCCCTGGATGAAAAAAGCCCACCTCGGCACAGGAACAGACAATGGTAATTTCTCTCCCGGCCTGGAGACCAAGAG
GCAACGGACAGCTTACACGAGACATCAGATCTTGGAGCTGGAGAAGGAGTTTCATTTCAACCGGTACCTTACAAGGCG
GAGACGGATAGAGATAGCTCACTTGCTATGTTTGTCAGAGAGGCAGATAAAAATCTGGTTCCAAAACCGGCGGATGAA
ATGGAAGAAAGACAACAAACTTCCTAACACGAAAAATGTGAAGAAAAGAAACCAACACGTAGAATCCCTGAACAACC
TGACACCCCTTTCGAACCACATACAACAACAACTAGGACGACTTCTTCCTCAGACAACACAACTAGACTCAAAATCTGA
CTACGGCTTAACAGAACTCTGATGTAACGTGAAATACGAAGAGGACCGCGTTGTGTACGGGATACGTGGGTTAATAAC
AGTCTTTCATGTAACAGGTCACTCTGTACTCGGCCAAATGTTCTGTGAACTTGTTTTATAGATGAAAAATTTGACCAACG
TCGTAGTTTAATTTAATTTGACATCCTGAATGAGTTGTGACATCATTTGTGTGTCTTTATTTTTTGTAAATATATTTAACC
AGGTCACTGTTGGGGTCATGCATTCAACAACTAAGGTTGACAAGTGAGCATTTTCTGCTTTATATATATATATTTGAGGCTC
TCACGTTGACTTGTATTACTCAGGGTATTTAGCTTTACACTTGTAGATTTAATTACTGACCGAGGAAAGAAGCTAGAGT
ATTCTTCATACTCCCATAAAATAAGAGAAACAGTGGTATTCTTTGTACTCCCATAAGATGTGAGAAACAGTGGTATTCT
TCATACTCCCATAAGATAAGAGAAACAGTGGTATTCTTTGTACTCCCATAAGATAAGAGAAACAGCAGTATTCTTCATA
CTTCCATAAGATAAGAGAAACAGTGGTATTCTTTGTACTCCCATAAGATGTGAGAAACAGCAGTATTCTTCATACTCCC
ATAAGATAAGAGAAACAGCAGTATTCTTCATACTCCCATAAGATAAGAGAAACAGCAGTATTCTTTGTACTCCCATAAG
ATAAGAGAAACAGCAGTATTCTTCATACTTCCATAAGATAAGAGAAACAGCAGTATTCTTCATACTCCCATAAGATAAG
AGAAACAGCAGTATTCTTCATACTCCCATAAGATAAGAGAAACAGCAGTGTTCTTCATACTCCCATAAGATAAGAGAA
ACGGCAGTATTGTTCATACTCCCATAAGATAAGAGAAACAGCAGTATTCTTTGTACTCCCATAAGATATGAGAAACAGC
AGTATTCTTCATACTCCCATAAGATAAGAGAAACAGCAGTATTCTTCATACTCCCATAAGATAAGAGAAACGGCAGTAT
TCTTCATACTCCCATAAGATAAGAGAAACGGCAGTATTCTTCATACTCCCGTATTTATATATTTTTTTTCTTGTCAACTGT
TACTACGAGAACCATTTAGCTCTGTAATAATGAGATCTTAATTTCGTGTCCGCTAAAGGTAAAATATGTAAAAATGATC
AAGATTATATGTTAAATATTATGTTTATTATAGTGTATGTATAGTATTTCCTTAATTTATATTCATTGCAGGTCATTACTT
TGATGAGTTAGCGTTGGAAAAGCAGTCCAGGTTCTTGTTCTAGACTTGTTTATTGCAGCTGACAGCGCCCCTTGTGGTTG
GAATTGAAAAATAAGATTTGGTTGTTTCCGTCATGGAACTGGTTGGTTGGAAGTAGATGTTGTGTTGGTTGTAATCTTCT
AGAAAGGATCAAACAAATATGCAAGAAAACATAATTAAAGGGCCATGTTTTGGCCATGTTGAAACGATACATTCCACG
AACTAGTACAGGCGACAGCAAAAAAATATGCTCTATTTTTATAGTCACGAGTCGGAGCTTTACAACACACTTTAAATTA
AAAAATAATATCATTTATTTTTAAGCCTAATTTCTAGTAACGTAATTAAAATATACGCTAGAACTAACTTGTTTAACTA
TCACAATGAAAGAAAATTGGTGTAAACCTTAAGTTATTTACAAGCGAATTTTCCATAAAATTTACAAAATTGAACTTAA
GCAACAAAGACCAACTTTTGAAACCAAAATTTAACATAAAGTCAACCTTTCTAATGAAGTTATATTCTCAAGCGAATCT
AATAATAGGTTTCTTAATTATCAAATGAAGTGTTTGATAAAAAATCAAAGTTTTCCACAAGTTACGTTCTTTTGTACAGA
TCCGGGTTCAGTCTTGAGTAGGTACGAGACATACATGCCACCAGTGTAAGCTGTATTGTTATTGGTCAACCAGGTCACG
TAGTCACTTTCCTCCGCACTTTTAGTTAACTCCTGATTTGTTTTTTAACAATTGTTGTGACCTTCTATTTAAAGATCTCTG
GCTGTATTGTTGTATAAGACAAATAAACACGCTTTGATTTGC

*Lpol-Dfd D*:

>Lpol_DfdD_XM_022387688.1 LOC106460884 [organism=Limulus polyphemus] [GeneID=106460884]
CATGTGATATGGTAATTTCTTTGGGAGGGCGGAGTCTAAGCCCTGTTAAACGACAAACGCGAAGAAAAATTATTTCCAT
CCAATTCACCCAAAAATTAATGATCATGAGCTCGTTTTTGATGAACTCACCGCCGTATTTGGAACCCAAATTTCCTCCCT
GCGAAGAATATTCACAGGTTAACTACATTCCTAACCAAGGCGAAAGCTATTACGCTCAGCAACCGCATCATCATCACG
GGTTTCACAATGCCGACACGCAGCACGCGCCGTTAAATTATGACCCTAACAGTGGCGGAGCCCGAGTTTCCGCTTTCAC
AAATCACGGAGCTACTAGTCCCTACTATCCGTCGTCATGTTCGCTGTCCCAGAGCAGTATATCAACACTTGAAAGGCTT
ACAAATCTGGGTTCCACCATAAATTGTGTAACGAGCGAACGTACTCGGGGCCGTGATGCTGTCTCTCCTGGACCAGGAC
AATCGCCCGAGAAGTCTCCACCGTCTTCGCCGCCATCTCCGCTGACAGATTCTAATTTCCACGCGGCGTCATCTTCTTCC
TCGCGGAATCAGCCGTCAACTAACGTTCTTTCGAGAGATTCACCTGTTCAGAATTCCTCTTCAAACCATGGAAACGAGT
CTCCCGATTTTGGGAACATTTCGAAACAACGACCGGCTATCTATCCGTGGATGAAGAAGGCTCATATGGGGTCGGCGA
ACGGCAACCTATCGCCGGGAATCGAGGCCAAACGGCAGAGGACGGCTTATACTAGACATCAAGTTCTTGAGTTAGAAA

AAGAGTTCCACTTTAATCGCTATTTAACACGACGTCGTCGAATTGAGATTGCTCATGCACTCTGCCTTTCAGAACGACA
GATTAAAATTTGGTTTCAGAACCGCCGTATGAAATGGAAGAAAGATAACAAGCTCCCTAACACCAAGAACGTCAGAAA
ACTGAGTCGCCATGCTGAAACCATTCACAGCGACCCATCGCCAGGTGTCCGACCCACTATCGGCCCGATCATGGGTCAC
CCATCATCACAAAACAGATCTGATCCAACGCGTCTAATCTCAAACCCCGAGAGCGGTGAAGTGTTACATTTGGGTTCCA
AAGGTAATTATGGATTAACGGAGCTCTGACTCGTCTAACTGAAAGAATAAAACATCTATAACTGAATGAAACCAGCAA
AAATTATCTGGACTCGTCTCACAGTCCAAGGGCGCAGGCATAAACTTAAAAGTCCAATCAATCCACATAACGCCTTTGA
GTATATAACTTATAGAATAATAGCCTAGGTTTATCCACTTGTGCAGATGTTTGTCTCTGGTATGGGAGAGTGAACACCT
AAGCTAAGGGAGAAAAACATATAAATTATCTACCAACACGAGATTAGGTTTAGGTGGTTAGAAGAACAACAAAAAAA
ATTGATGTTATCGAGTATCCACGTACGTTG

*Lpol-Dfd E*:

>Lpol_DfdE_XM_022379496.1 LOC106474221 [organism=Limulus polyphemus] [GeneID=106474221]
CATGTGATATGGTAATTTCCTCAAAAGGGCGGAGTTTAAGCCCTGTTAAACGACATACGCGAACGAAAATTATTTCCAT
CCAACTCATTCAAAAATTAATGATCATGAGCTCGTTTTTGATGAACTCACAACCGTTTATGGAGCCAAAATTTCCACCTT
GCGAAGAATATGCACAAGCTAGCTACATTCCATGCCATGGCACAAGCTATTTCTCTCCACAAGCGCAGCATCATGACG
AGTTCCATAGCGCTGACACTCAGCACGCGCCGATAAATTATGACCTTAATACCGGTGGAGCTAGAACAGCCCCTTACAC
AAATCACGGTCCTGCTAACTCTGTCTTTCCGGCATCATGTTCAATGTCTCAGAACAGTATATCAACACTTGAAAAGCTTA
CAAATCTAGGCTCCTCCATAAATTGTGTAACGAGCGTACGAGTTCGGAAAATTAATGCTACTCCCCACGGACTAGGACA
ATCTCCTTATAGCAGTGATGCATCTCCTGAGTCCTCTCCACCACCACCAGAAGATACTTTACAAACAGCACCAGTTCCTT
CTCTACAAGTCCAACCATTAACCAGTATTCACAATAGAGATTCGTCGGTCAAGAATTCCTCTTCAATTCTGAGAAGCGA
GTCTCCAGAATGTGTAAATTCCCCTGAAGTACAACCGACAATCTACCCCTGGATGAAGAAAGTTCATCTTGGGTCAGCA
AATGGAAGTTGTTTTTCATCAGGGATAGAAACTAAGCGCCAGCGGACAGCCTATACTAGACATCAGGTTCTGGAGTTA
GAGAAAGAGTTCCACTTTAACCGTTATTTGACACGACGTCGCCGAATTGAGATTGCCCATGCACTCTGCCTTTCAGAGC
GTCAGATTAAAATTTGGTTTCAGAACCGTCGTATGAAGTGGAAGAAAGACAACAAGCTACCGAATACAAAAAATGTGA
GAAAAATTAGTCGGCAGGTCGAGAACCTTCACCACGGTCTCTCAACATCGCCCGGTGTTAGACCCACTCTCAGTCCGCT
TGCGAGTCACCTTCCTTCGCGGCAAGATAGACGCACGGTGGATCCAATAAATATAACAAATCCCTAAAACAGTGAAAT
ACTACATACTGGATGTAAAAGTCACTATGGATTAACAGAGCTTTGACCCAATCAATGACAAAGAGTGTGATCGTTAGAT
GTTTGAGTTCTGCAACTAATTTTGTGGCTCATTGATTTACTCTACAACTCACTGGCGTTTGTACGTAATTAATATCCATA
ATCCTTTCTGTCTCAACACTTTGATTTTGCTACCAGAGACTATAAGGAGATTCTACTAATCTGTGCACTTCGCTGGTAAG
TGAAGGTAATCTTCTAAGAGGGAGTATCAAAACGACGGAACCAAGAATCTTAAAAATGAGAAAATTATACTTGAATAT
AAATACAGTGAGGAATATCTTCGCGCTTTTATAACTTAATTTATAAGTTTATGAATATGTTATTATTCAAATGTAAGG

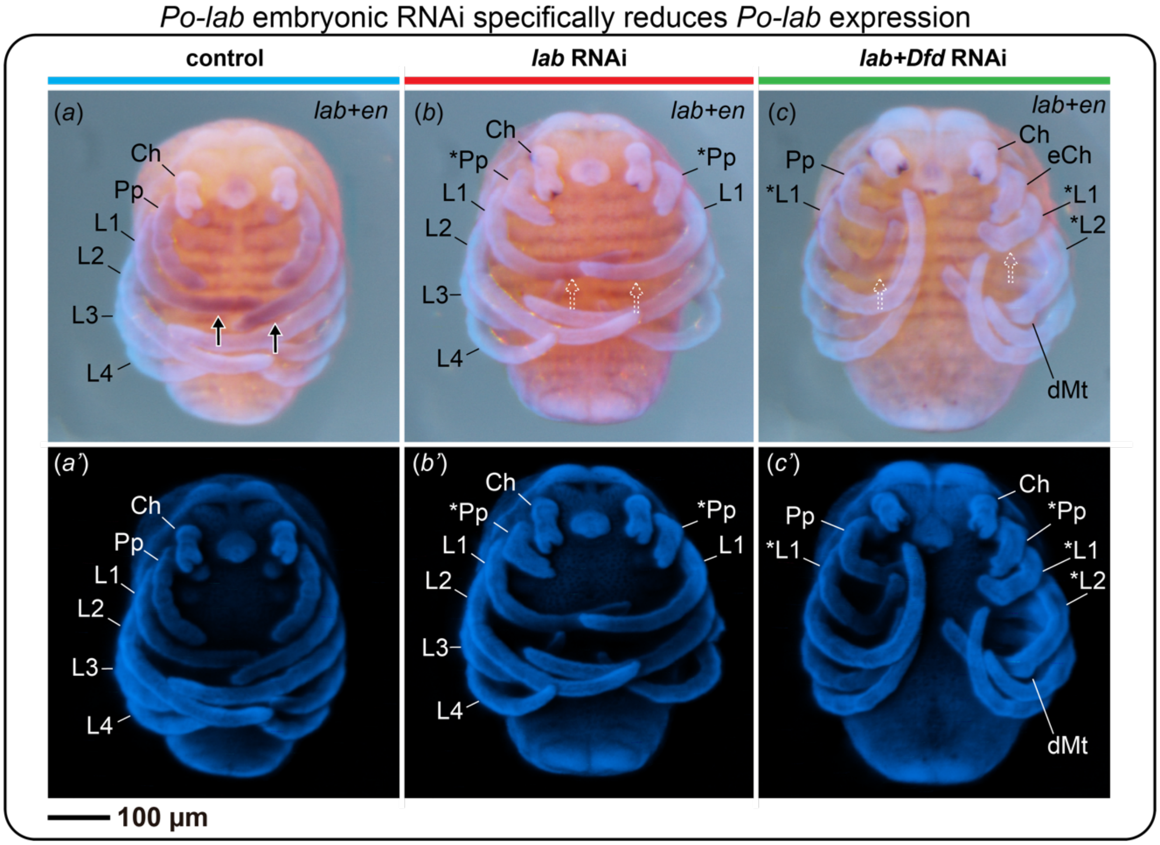


**Figure S1**. Double colorimetric in situ hybridization of *Po-en* and *Po-lab* in stage 11 embryos. Bright field images are merged with nuclear counterstaining (Hoechst). a: control. b: *Po-lab* RNAi. c: *Po-Dfd* RNAi. a'-c': Corresponding embryos counterstained for nuclei (Hoechst). Arrows: strong Po-lab expression on distal leg 2. Dotted arrow: attenuated expression on distal leg 2. Asterisks: homeotic appendages mark. Ch: chelicera; dMt: defective metatarsus. L1–4: legs 1–4; Pp: pedipalp. Scale bar: 100 $\mu$m.

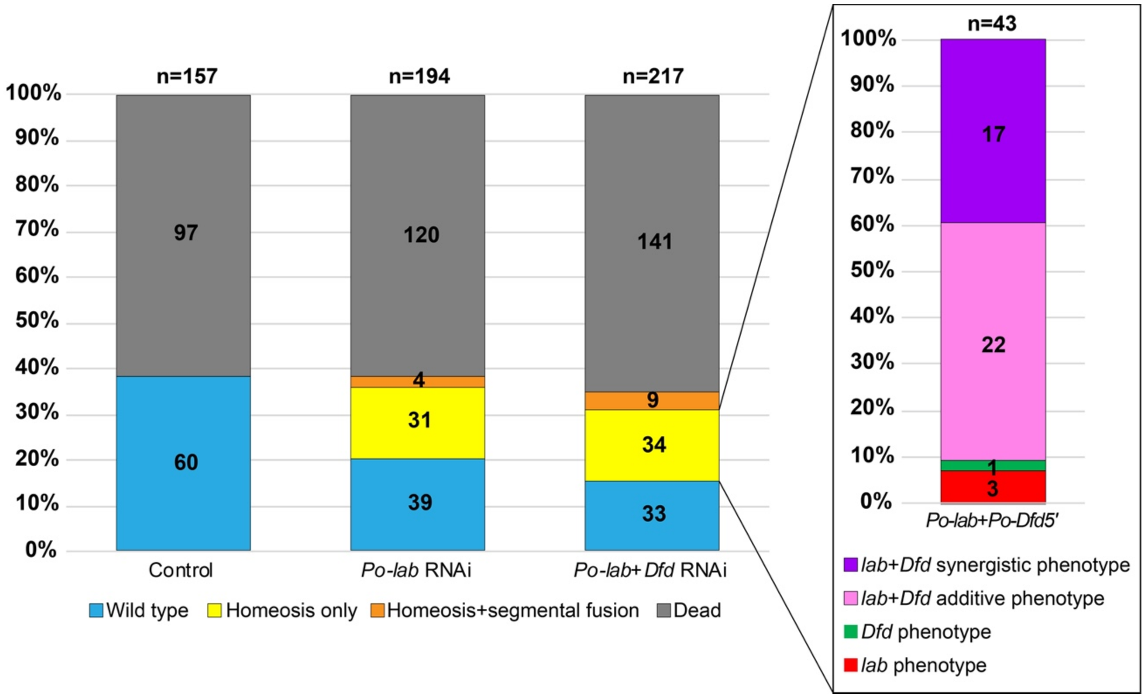


**Figure S2.** Quantification of phenotypic classes in control (left), *Po-lab* RNAi (center) and double knockdown *Po-lab* + *Po-Dfd* RNAi (right). Box: Phenotypic breakdown of the double knockdown individuals exhibiting a *Po-lab* + *Po-Dfd* phenotype.

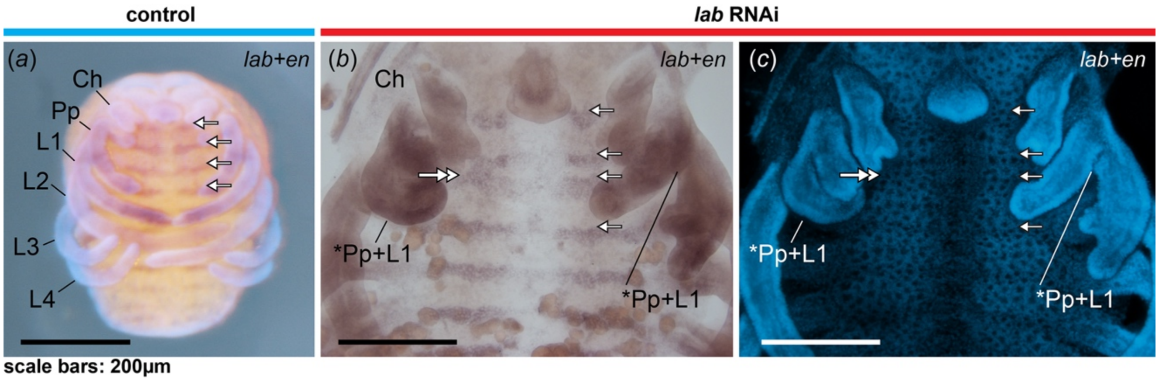


**Figure S3.** Segmental fusion upon *Po-lab* knockdown. Double colorimetric in situ hybridization for *Po-lab* and *Po-en*. (a) Control embryo. Hoechst nuclear stained overlaid with brightfield image. (b) *Po-lab* RNAi embryo. Brightfield image. Note fusion of engrailed stripes on the left side (stronger transformation) and reduced spacing of adjacent stripes on the right (weaker transformation). (c) Nuclear counter stanning. Arrows: *Po-en* expression stripes marking posterior boundaries of segments. Asterisks: homeotic appendages mark. Ch: chelicera; L1–4: legs 1–4; Pp: pedipalp. Scale bar: 200 $\mu$m.

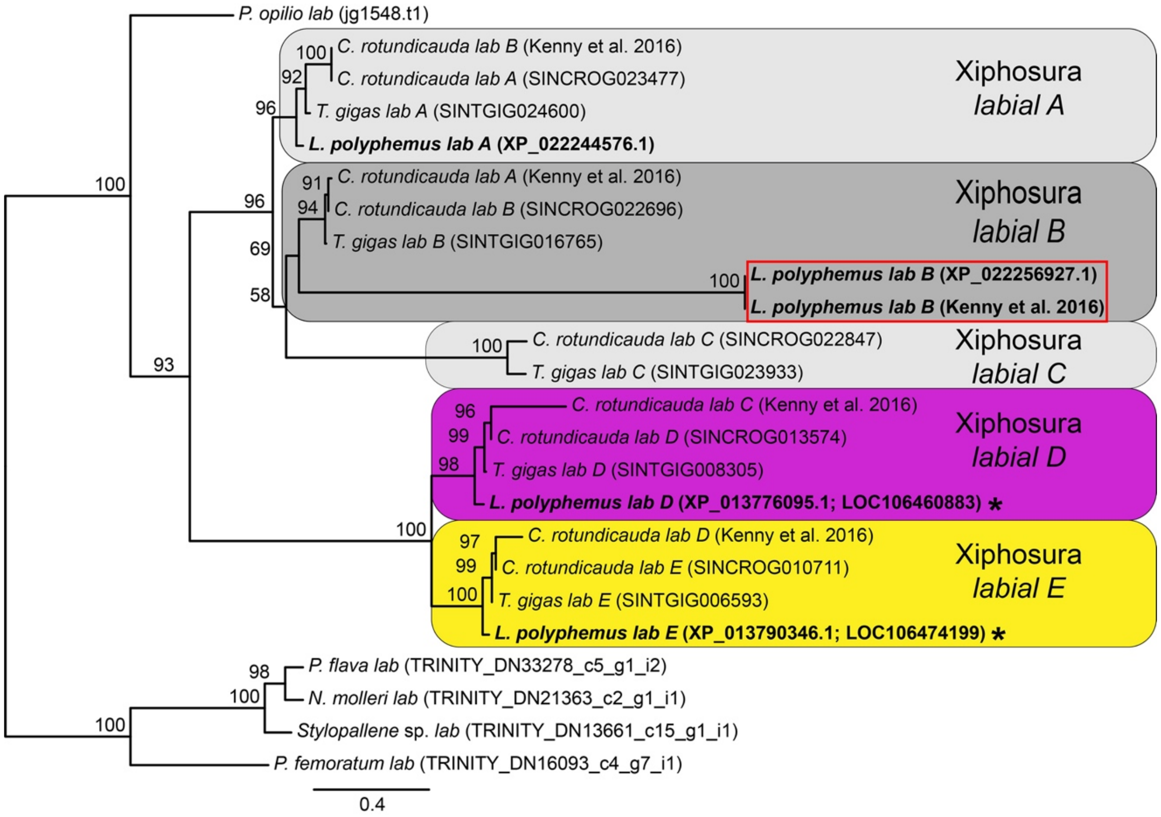


**Figure S4.** Maximum likelihood phylogenetic analysis (IQTREE) of *labial* homologs in horseshoe crabs and selected outgroups. Numbers in the nodes are ultrafast bootstraps. Red box indicates *L. polyphemus* unusual *labial* copy (see Supplementary Methods). Asterisks marks *L. polyphemus* paralogs assayed with HCR.

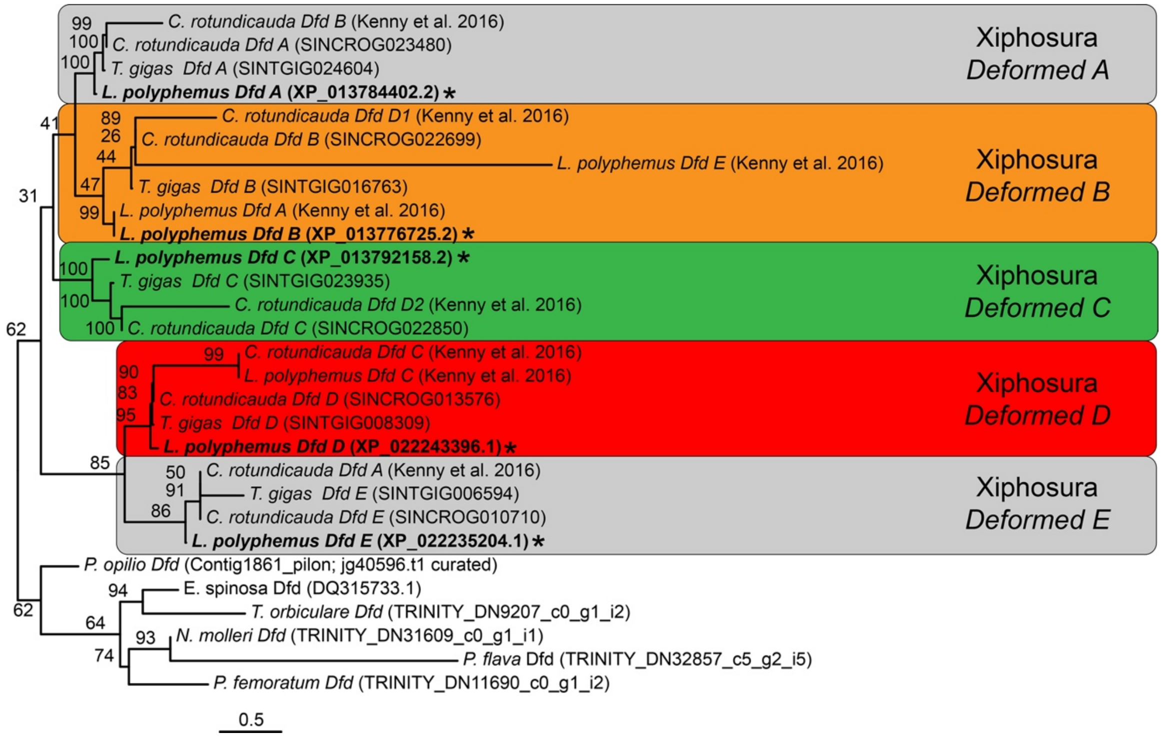


**Figure S5.** Maximum likelihood phylogenetic analysis (IQTREE) of *Deformed* homologs in horseshoe crabs and selected outgroups. Numbers in the nodes are ultrafast bootstraps. Asterisks marks *L. polyphemus* paralogs assayed with HCR.

**Table S1.** Accession numbers and primers for the genes studied in *Phalangium opilio*.

| Gene ID | Gene | Primer ID | Forward | Reverse | Product size (bp) | Comments |
|---|---|---|---|---|---|---|
| jg1548.t1 | *Po-labial* | Popi_lab | GAACCCGATTGGTTGGATAA | CGGACTGTCTCCTCCTCAAG | 746 | Sharma et al. 2012 |
| jg1352.t1 | *Po-proboscipedia* | Popi_pb | TCCAAAATCGGAGGATGAAG | CGAAGACGAAGGTGAAGAGG | 998 | Sharma et al. 2012 |
| jg40596.t1 | *Po-Deformed* | Popi_Dfd_5' | ggccgcggTTTCTGCCGCTACGACTTTG | cccggggcTACGGCGGAGAGTTCATCAA | 812 | Gainett et al. 2021 |
| jg16360.t1; Popi1_comp41585_c0_seq1; Popi2_comp171093_c0_seq2 | *Popi-engrailed* | Popi_en | ggccgcggCGTCCGATTTTTACGTTCTCA | cccggggcCGTTAACTCCTCCGTTAGGC | 718 | Sharma et al. 2012 |

**Table S2.** Accession number of *labial* and *Deformed* homologs used in the phylogenetic analyses.

| **Species** | **Gene** | **Gene ID** | **Database** | **Reference** |
|---|---|---|---|---|
| *Carcinoscorpius rotundicauda* | *Crot-labial A* | SINCROG023477 | GCA_011833715.1 | Shingate et al. 2020a |
| *Carcinoscorpius rotundicauda* | *Crot-labial B* | SINCROG022696 | GCA_011833715.2 | Shingate et al. 2020a |
| *Carcinoscorpius rotundicauda* | *Crot-labial C* | SINCROG022847 | GCA_011833715.3 | Shingate et al. 2020a |
| *Carcinoscorpius rotundicauda* | *Crot-labial D* | SINCROG013574 | GCA_011833715.4 | Shingate et al. 2020a |
| *Carcinoscorpius rotundicauda* | *Crot-labial E* | SINCROG010711 | GCA_011833715.5 | Shingate et al. 2020a |
| *Limulus polyphemus* | *Lpol-labial A* | XP_022244576.1; LOC111086353 | GCF_000517525.1 | Battelle et al. 2015 |
| *Limulus polyphemus* | *Lpol-labial B* | XP_022256927.1; LOC106472664 | GCF_000517525.1 | Battelle et al. 2015 |
| *Limulus polyphemus* | *Lpol-labial D* | XP_013776095.1; LOC106460883 | GCF_000517525.1 | Battelle et al. 2015 |
| *Limulus polyphemus* | *Lpol-labial E* | XP_013790346.1; LOC106474199 | GCF_000517525.1 | Battelle et al. 2015 |
| *Tachypleus gigas* | *Tgig-labial A* | SINTGIG024600 | GCA_014155125.1 | Shingate et al. 2020b |
| *Tachypleus gigas* | *Tgig-labial B* | SINTGIG016765 | GCA_014155125.1 | Shingate et al. 2020b |
| *Tachypleus gigas* | *Tgig-labial C* | SINTGIG023933 | GCA_014155125.1 | Shingate et al. 2020b |
| *Tachypleus gigas* | *Tgig-labial D* | SINTGIG008305 | GCA_014155125.1 | Shingate et al. 2020b |
| *Tachypleus gigas* | *Tgig-labial E* | SINTGIG006593 | GCA_014155125.1 | Shingate et al. 2020b |
| *Nymphon molleri* | *Nmol-lab* | TRINITY_DN21363_c2_g1 | | Ballesteros et al. 2020 |
| *Pallenella flava* | *Pfla-lab* | TRINITY_DN33278_c5_g1 | | Ballesteros et al. 2020 |
| *Phoxichilidium femoratum* | *Pfem-lab* | TRINITY_DN16093_c4_g7 | | Ballesteros et al. 2020 |
| *Stylopallene cheilorhynchus* | *Sche-lab* | TRINITY_DN13661_c15_g1 | | Ballesteros et al. 2020 |
| *Carcinoscorpius rotundicauda* | *Crot-Dfd A* | SINCROG023480 | GCA_011833715.1 | Shingate et al. 2020a |
| *Carcinoscorpius rotundicauda* | *Crot-Dfd B* | SINCROG022699 | GCA_011833715.2 | Shingate et al. 2020a |
| *Carcinoscorpius rotundicauda* | *Crot-Dfd C* | SINCROG022850 | GCA_011833715.3 | Shingate et al. 2020a |
| *Carcinoscorpius rotundicauda* | *Crot-Dfd D* | SINCROG013576 | GCA_011833715.4 | Shingate et al. 2020a |
| *Carcinoscorpius rotundicauda* | *Crot-Dfd E* | SINCROG010710 | GCA_011833715.5 | Shingate et al. 2020a |
| *Limulus polyphemus* | *Lpol-Dfd A* | XP_013784402.2; LOC106468515 | GCF_000517525.1 | Battelle et al. 2015 |
| *Limulus polyphemus* | *Lpol-Dfd B* | XP_013776725.2; LOC106461448 | GCF_000517525.1 | Battelle et al. 2015 |
| *Limulus polyphemus* | *Lpol-Dfd C* | XP_013792158.2; LOC106476035 | GCF_000517525.1 | Battelle et al. 2015 |
| *Limulus polyphemus* | *Lpol-Dfd D* | XP_022243396.1; LOC106460884 | GCF_000517525.1 | Battelle et al. 2015 |
| *Limulus polyphemus* | *Lpol-Dfd E* | XP_022235204.1; LOC106474221 | GCF_000517525.1 | Battelle et al. 2015 |
| *Tachypleus gigas* | *Tgig-Dfd A* | SINTGIG024604 | GCA_014155125.1 | Shingate et al. 2020b |
| *Tachypleus gigas* | *Tgig-Dfd B* | SINTGIG016763 | GCA_014155125.1 | Shingate et al. 2020b |
| *Tachypleus gigas* | *Tgig-Dfd C* | SINTGIG023935 | GCA_014155125.1 | Shingate et al. 2020b |
| *Tachypleus gigas* | *Tgig-Dfd D* | SINTGIG008309 | GCA_014155125.1 | Shingate et al. 2020b |

| | | | | |
|---|---|---|---|---|
| *Tachypleus gigas* | *Tgig-Dfd E* | SINTGIG006594 | GCA_014155125.1 | Shingate et al. 2020b |
| *Endeis spinosa* | *Espi-Dfd* | DQ315733.1 | | Ballesteros et al. 2020 |
| *Pallenella flava* | *Pfla-Dfd* | TRINITY_DN32857_c5_g2_i5 | | Ballesteros et al. 2020 |
| *Nymphon molleri* | *Nmol_Dfd* | TRINITY_DN31609_c0_g1_i1 | | Ballesteros et al. 2020 |
| *Phoxichilidium femoratum* | *Pfem-Dfd* | TRINITY_DN11690_c0_g1 | | Ballesteros et al. 2020 |
| *Tanystylum orbiculare* | *Torb-Dfd* | TRINITY_DN9207_c0_g1_i2 | | Ballesteros et al. 2020 |

**Table S3.** List of plasmids and clones for the genes used in RNAi experiments in *Phalangium opilio*.

| Genome ID | Gene | Primer | Plasmid # | Plasmid ID | Direction | Sequence (with vectors endings) |
|---|---|---|---|---|---|---|
| jg1548.t1 (curated); Contig9232_pilon | *Popi-labial* | *Popi_lab* | 2 | BTX681 | Forward | AAAGATCTTAGGGCGAATTGAATTTAGCGGCCGCGAATTCGCC<br>CTTCGGACTGTCTCCTCCTCAAGCCGCGTCCGTACCAACTTATA<br>AATGGATGCAAGTGAAAAGAAATGTTCCAAAACCAGTCAACAA<br>AACAGAATTCGGTTTCAGCGGCGGAGGAAACATGGTGGGTAGC<br>GGAGGTAACGGCGGTGGCTTGGGCGGAATCGCGGGTACGGGA<br>ATGTGCGGTGGAGCCGGCGGCGGTCTAAACGGATCGAACCTCG<br>GCAACGGTTTAGCCGGTGGTCCCGGTTCGGGTCGGACAAACTTT<br>ACGACGAAACAACTAACCGAACTCGAAAAAGAATTTCACTTTA<br>ACAAGTACCTGACCAGGGCCAGGCGAATCGAAATCGCCACCGC<br>CTTGCAACTGAACGAGACGCAGGTCAAAATATGGTTTCAAAAT<br>CGTCGAATGAAACAAAAGAAACGTATGAAAGAGGGTTTAATAC<br>CGCCGGAACCCATTTCAACGGACAGTTCTTCTCCCCCGTCATCC<br>GCCCAATCTCCCGGTCAACCAAACTCAGTCGGCGGTAATGGAG<br>CTTTACTCGCCGGTGGTAGTGATAACAGCCAGTGACCGGTCCGC<br>TTCTCACCCCAAAATCGCGATATTGTCATCGAGTCACCAATTGT<br>AAATAAATAAACAACTAACAAGATACGAAACACTAAACAACTT<br>CGGATGGTTACTGGAAAAAAAAAATTAGAAAACTATCTCCATT<br>CAGTGATTACTCCAACATATAAACAATAAAA |
| jg1548.t1 (curated); Contig9232_pilon | *Popi-labial* | *Popi_lab* | 3 | BTX682 | Forward | AACGATCTTAGGGCGAATTGAATTTAGCGGCCGCGAATTCGCC<br>CTTCGGACTGTCTCCTCCTCAAGCCGCGTCCGTACCAACTTATA<br>AATGGATGCAAGTGAAAAGAAATGTTCCAAGACCAGTCAACAA<br>AACAGAATTCGGTTTCAGCGGCGGAGGAAATATGGTCGGTAGC<br>GGAGGTAACGGCGGTGGCTTGGGCGGAATCGCGGGTACGGGA<br>ATGTGCGGTGGAGCCGGCGGCGGTCTAAACGGATCGAATCTCG<br>GCAACGGTTTAGCCGGTGGTCCCGGTTCGGGTCGGACAAACTTT<br>ACGACGAAACAACTAACTGAACTCGAAAAAGAATTTCACTTTA<br>ACAAGTACCTGACCAGGGCCAGGCGAATCGAAATCGCCACCGC<br>CTTGCAATTGAACGAGACGCAGGTCAAAATATGGTTTCAAAAT<br>CGTCGAATGAAACAAAAGAAACGTATGAAAGAGGGTTTAATAC<br>CGCCGGAACCCATTTCAACGGACAGTTCTTCTCCCCCGTCATCC<br>GCCCAATCTCCCGGTCAACCAAACTCAGTCGGCGGTAATGGAG<br>CTTTACTCGCCGGTGGTAGTGATAACAGCCAGTGACCGGTCCGC<br>TTCTCACCCCAAAATCGCGATATAATGTCATCGAGTCACCAATT<br>GTAAATAAATATACAACAAACAAGAAACGAAACACTAAACAA<br>CATCGGATGGTTAGTGGAAAAAAAAAATTAAAAAAACAATTTC<br>AAATTCAGTGATTACGTCAAACAAAGAAACAATAAAAGACTAA<br>AATTATCCAACCAATCGGGTTCAAGGGCGAATTCGTTTAAACCT<br>GCAGGACTAGTCCCTTTAGTGAGGGTTAATTCTGAGCTTGGCGT<br>AATCATGGTCATAGCTGTTTCCTGTGTGAAATTGTTATCCGCTC<br>ACAATTCCACACAACATACGAGCCGGAAGCATAAAGTGTAAAG<br>CCTGGGGTGCCTAATGAGTGAGCTAACTCACATTAATTGCGTTG<br>CGCTCACTGCCCGCTTTCCAGTCGGGAAACCTGTCGTGCCAGCT<br>GCATTAATGAATCGGCCACGCGCGGGGAGAGGCGTTTGCGTAT<br>GGGCGCTCTTCCGCTCCTCGTTCCTGACTCGCTGGCTCGGTCGT<br>TCGGTNCGGNAGGGTATCAGTCCCTCAAGGCGG |
| jg40596.t1 (curated); Contig1861_pilon | *Popi-Deformed* | *Popi_Dfd_5'* | 1 | AXW413 | Reverse | NNNNNNNNNNNNNNNNNNTNANCNNNNNTNNNGGACTAGTCCT<br>GCAGGTTTAAACGAATTCGCCCTTCCCGGGGCTACGNCGNNNN<br>GTTCATCAAAAACGAACTCATGATCATTAATTTTTGGGTGAATT<br>GGATGGAAATAATTTTTCTTCGGGGTTGTCGTTTAACCGCGCGC<br>TGGCTCCGCCTCTCGGCAGAAATTAGCATATCACGCGGCCGCC<br>GTCGTCCAATCGGCTCGGTGGGCTTTGACCCGGGGCGCTCGCGT<br>CAGATTGATCGATGACAGTGACACATTGCGAGAAAATGGCGGC<br>GCCGGCCATACCTGTAGGCTAGGCCCCGCCGCCCCGCCAACGG<br>ACGCGTGAAAATATATTTTTCAAACACAATATACTCGAAAACG<br>CGCTCGTCGCGCGCTCTAGGATCTCGCGCCGCCGCCGAAATTCA<br>CCACAACAGTTCCTCGTGCAAAATCCCCGTTCTTGAGTGGAAAT<br>GCGATGCAAAAATGATGCCTACCGTAATGTCCTCAGGCTCGAA<br>CACCAACAATCAAGCGTGCAAAATGCTTTATCAGCCGATGTAC<br>TCGGTCCTTAAGGGAGAACTCGGGAAGACTCCGAGCGATTGGA<br>ACAACTACGGAGATTGTTGTTGGGTCGAGCTAAATTGAAAGCT<br>CTTTCGTTTCGATTAAAAAAAAAGGAGCCCGTTGAAGAAGGGCT<br>CGGACGAGTCGTTAAGAGCGCTTGATACGACTCGTGGTGGAGG<br>TTTGAAACGGCAGCCGCGAAGTGCAACTTCTAACGTGGATATG<br>ACGTTAAGAGCCGANTTTCGCGCTAACCCACACCAACTATGTCT<br>GATCAACGTGTGCCAGTAGCAACGTGCACAAAGTCGTAGCGGG<br>CAGAAACCGCGGCCAAGGGCGATTNNGCGGCCGCTAAATTNNA<br>ATTCGCCCNTATAGTGAGTCGTATTNNCATTCNNCTGGGCCGTC<br>GTTTTANACGTCNNGACNNGGNNAAANNTGGNNGTTACNNACT<br>TANNCNCNNGCAGCCACATTCCCCTTNNGCNNNCNNGGNNNNA<br>TNNCNANAGGNCNNNNNCGANNNCCCNNNNCNAACNNTTNNN<br>NNNCNNNNNCNNNNCNGNANTNNNNNNNNCNNNNNNNNNNA<br>NNTNNNNNNNTNNNNNNNNNTTGNNGNNNNGTNNNNNNNNNG<br>GNANNNNTNN |

| | | | | | | |
|---|---|---|---|---|---|---|
| jg40596.t1 (curated); Contig1861_pilon | *Popi-Deformed* | *Popi_Dfd_5'* | 2 | AXW414 | Reverse | NNNNNNNNNNNNNNNNNNNNANNNNCACTNNNGGACTAGTCCTGCAGGTTTAAACGAATTCGCCCTTCCCGGGGCTANNNCNNANAGTTCATCAAAAACGAACTCATGATCATTAATTTTTGGGTGAATTGGATGGAAATAATTTTTCTTCGGGGTTGTCGTTTAACCGCGCGCTGGCTCCGCCTCTCGGCAGAAATTAGCATATCACGCGGCCGCCGTCGTCCAATCGGCTCGGTGGGCTTTGACCCGGAGCGCTCGCGTCAGATTGATCGATGACAGTGACACATTGCGAGAAAATGGCGGCGCCGGCCATACCTGTAGGCTAGGCCCCGCCGCCCCGCCAACGGACGCGTGAAAATATATTTTTCAAACACAATATACTCGAAAACGCGCTCGTCGCGCGCTCTAGGATCTCGCGCCGCTGCCGAAATTCACCACAACAGTTCCTCGTGCAAAATCCCCGTTCTTGAGTGGAAATGCGATGCAAAAATGATGCCTACCGTAATGTCCTCAGGCTCGAACACCAACAAACGAGCGTGCAAAATGCTTTATCAGCCGATGTACTCGGTCCTTAAGGGAGAACTCCGAGCGATTGGAGCAACTACGGAGATTGTTGTTGGGTCGAGCTAAATTGAAAGCTCTTTCGTTTCGATTAAAAAAGGAGCCCGTTGAGGAAGGGCTCGGACGAGTCGTTAAGAGCGCTTGATGCCGTTCAACGTTGCTATACGACTCGTGGTGGAGGTTTGAAACGGCAGCCGCGAAGTGCAACTTCTAACGTGGATATGACGTTAAGAGCCGATTTCGCGCTAACCCACACCAACTATGTCTGATCAACGTGTGCCAGTAGCAACGTGCACAAAGTCGTAGCGGCAGAAACCGCGGCCAAGGGCGAATTCGCGGCCGCTAAATTCAATTCGCCCTATAGTGAGTCGTATTACAATTCACTGGCCGTCGTTTTACAACGTCGTGACTGGGAAAACCCTGGNGTTACCCAACTTAATCGCCTTGCAGCNNATCCCCCTTTCGCCAGCTNNNNNTANNANCNAAGAGGCCCGCACCGATCGCCCTTCCCAACAGTTGGCCAGCCTNNACGTACGGNNGGTTTAAGGTTTANCCNNAAAAANGAAAGCCGTTNNNNNNGGNTTT |
| jg1352.t1 (curated); Contig3674_pilon | *Popi-proboscipedia* | *Popi_pb* | 1 | BTX677 | Reverse | TAAGATCTTAGGGCGAATTGAATTTAGCGGCCGCGAATTCGCCCTTCGAAGACGAAGGTGAAGAGGACGTTTTATAACTCGCTCCGGATAGACCGTAACTCGTGTCCATGTGTTGCGATTGAATGACCGACTGTTGCTGATGAGTTGTCGGTGGTGAAGTGCGGTAACAATAAGGTGGGGCCATGGTTGGCGAGGATGTTCGGTGGTTTTGAGTCGAAAAGTAAGACGACGCGGAACTAGCGTCCGAATGGTGGTAATTGTTGGCGTAAAGCCTCGACCTTTGGTAACTGGGCGAGGCCGACTGCATCCTGCTTTGCAGGGCGGCGTGTGCCTGAGAAGGCGTTTGGTAGGAACCTCCCGGACTGGCCGGTTGTCCCGCAGTCCCGCACTGAACCGGGTTACAAGGCTGATGACATCTGCTCGTCCCGTTCATATCGATGATACATGACAACGACTTTGTCGTGCTTGGAGAACCGACAGAAGAAAGACTATTTCTGCTAGATGGTGATCCGTTTAGACGACCGCTCAAGGCCTCGCTAACCCCGCCCGCCGTTAGGATGGTTGTTTTACCGCCGCTAAGAGGATTCAAAGTTCTTTCGACTCCGCTTAACGAAGATGAGAGCAAACCTTCCGAAACGGAACTGAGGTCCATCGTCCGATCCCCGTCGTCGGCGACGAAATGGAGTTGCCGGAACACGTCGTCGGAGTGGCGGTTGAGCTAGGTACGTGCGTCGATGGCGAATGAAGATGTCTCATAGAATTCTCGCCCTCGGACATGAGAATCGAATCTGATTTTGCCGGGGAAACCGTCCTGCCGCAACATAGATCCGATTTGGATCCGCCACCGCCCATGGGCGTCAAAGGCGTCGATCCATCCTGGCTGGTAGAAGTCTGATCGGAACAGGGACTCCCGATCGACGACAGGGTGGACGGACCTTTGTCCGACATGGAATCTTTACCATTCTCCGACTGTCCGCTACCCTTTTCATCGCCGTTCTTAGTCATCAAAGACTGTCTCTTGTGCTTCATCCTCCGATTTTGGAAAGGGCGAATTCGTTTAACCTGCAGGACTAGTCCCTTTATGGAGGTTAATTCTGAGCTTGGCTAATCTTGGGCANAGCTGTTTCCTGGGGNAAA |
